# FXa cleaves the SARS-CoV-2 spike protein and blocks cell entry to protect against infection with inferior effects in B.1.1.7 variant

**DOI:** 10.1101/2021.06.07.447437

**Authors:** Wenjuan Dong, Jing Wang, Lei Tian, Jianying Zhang, Heather Mead, Sierra A. Jaramillo, Aimin Li, Ross E. Zumwalt, Sean P.J. Whelan, Erik W. Settles, Paul S. Keim, Bridget Marie Barker, Michael A. Caligiuri, Jianhua Yu

**Affiliations:** Department of Hematology & Hematopoietic Cell Transplantation, City of Hope National Medical Center, Los Angeles, CA 91010, USA; Hematologic Malignancies Research Institute, City of Hope National Medical Center, Los Angeles, CA 91010, USA; Department of Computational and Quantitative Medicine, City of Hope National Medical Center, Los Angeles, CA 91010, USA; Pathogen and Microbiome Institute, Northern Arizona University, Flagstaff, AZ 86011, USA; Pathology Core of Shared Resources Core, Beckman Research Institute, City of Hope National Medical Center, Los Angeles, CA 91010, USA; Department of Pathology, University of New Mexico, Albuquerque, NM 87131, USA; Department of Molecular Microbiology, Washington University School of Medicine, St. Louis, MO 63110, USA; City of Hope Comprehensive Cancer Center, Los Angeles, CA 91010, USA; Department of Immuno-Oncology, City of Hope, Los Angeles, CA 91010, USA

## Abstract

The ongoing coronavirus disease 2019 (COVID-19) pandemic is caused by infection with severe acute respiratory syndrome coronavirus 2 (SARS-CoV-2). Human natural defense mechanisms against SARS-CoV-2 are largely unknown. Serine proteases (SPs) including furin and TMPRSS2 cleave SARS-CoV-2 spike protein, facilitating viral entry. Here, we show that FXa, a SP for blood coagulation, is upregulated in COVID-19 patients compared to non-COVID-19 donors and exerts anti-viral activity. Mechanistically, FXa cleaves the SARS-CoV-2 spike protein, which prevents its binding to ACE2, and thus blocks viral entry. Furthermore, the variant B.1.1.7 with several mutations is dramatically resistant to the anti-viral effect of FXa compared to wild-type SARA-CoV-2 *in vivo* and *in vitro*. The anti-coagulant rivaroxaban directly inhibits FXa and facilitates viral entry, whereas the indirect inhibitor fondaparinux does not. In a lethal humanized hACE2 mouse model of SARS-CoV-2, FXa prolonged survival while combination with rivaroxaban but not fondaparinux abrogated this protection. These preclinical results identify a previously unknown SP function and associated anti-viral host defense mechanism and suggest caution in considering direct inhibitors for prevention or treatment of thrombotic complications in COVID-19 patients.

## Introduction

SARS-CoV-2 is the pathogen responsible for the global COVID-19 pandemic^1^. To date, over 170,000,000 cases and approximately 3,500,000 deaths have been recorded with a worldwide mortality rate of 2%^2^. The public health and economic consequences have been devastating. Although some strategies such as neutralizing antibodies and vaccines are being developed or testing in the clinic, the pandemic rages on with new concerns regarding the emergence of resistant strains^3–6^.

Angiotensin-converting enzyme 2 (ACE2) has been identified as the host receptor for SARS-CoV-2^7,8^. SARS-CoV-2 uses its spike (S) protein to bind to ACE2 and enter host cells. Several host serine proteases (SPs) have been identified as facilitating SARS-CoV-2 entry via cleavage of its S protein into functional S1 and S2 subunits^9,10^. Furin cuts the S protein at the PRRAR (R-R-A-R685 ↓) site into S1 and S2 subunits at virus budding, while TMPRSS2 cleaves at the S2’ site (P-S-K-R815 ↓) at virus entry, and both cleavages enhance efficiency of SARS-CoV-2 infection^9,11^. Another SP family member, coagulation factor Xa (FXa) binds to tissue factor to initiate conversion of prothrombin to thrombin in the clotting cascade^12^. The direct FXa inhibitors rivaroxaban, apixaban, and edoxaban, and betrixaban as well as the indirect inhibitor fondaparinux have been developed as clinical anti-coagulants^13^, and several direct inhibitors are currently being evaluated for use in patients at high-risk for COVID-19^14^.

Here we show that FXa inhibits SARS-CoV-2 entry. Mechanistically, FXa binds to and cleaves S protein, which produced a different cleavage pattern than that of furin and TMPRSS2, and blocked S protein binding to ACE2. The effect was pronounced for the ancestral wild type variant, but was diminished in the B.1.1.7 variant. Exogenous FXa protected mice from lethal infection in a humanized hACE2 mouse model of COVID-19 using the wild type variant but not the B.1.1.7 variant. The anti-viral effect of FXa was attenuated by the direct FXa inhibitor rivaroxaban (RIVA) but not the indirect inhibitor fondaparinux (FONDA) both *in vivo* and *in vitro*.

## Results

To identify changes in SPs during SARS-CoV-2 infection, we examined their expression in lung samples from COVID-19 patients using an immunohistochemistry assay (IHC). Due to the lack of specific antibodies directly against FXa, we detected FX expression since ~100% FX can be activated to FXa at injury sites when platelets are exposed to both collagen and thrombin^15^. Our IHC analysis indicated that thrombin was highly expressed in lungs from COVID-19 patients compared to that from non-COVID-19 patients (Extended Data Fig. 1a), consistent with previous data^16,17^. We found FX was significantly increased in the lungs of COVID-19 patients compared to non-COVID-19 donors; in contrast, we did not observe consistent upregulation of other tested SPs (Fig. 1a and Extended Data Fig. 1b). We found that FX was also increased in the liver (Extended Data Fig. 1c) and serum, which is the source and carrier of FXa, in COVID-19 patients compared to non-COVID-19 donors (Fig. 1, b and c).

**Fig. 1.**
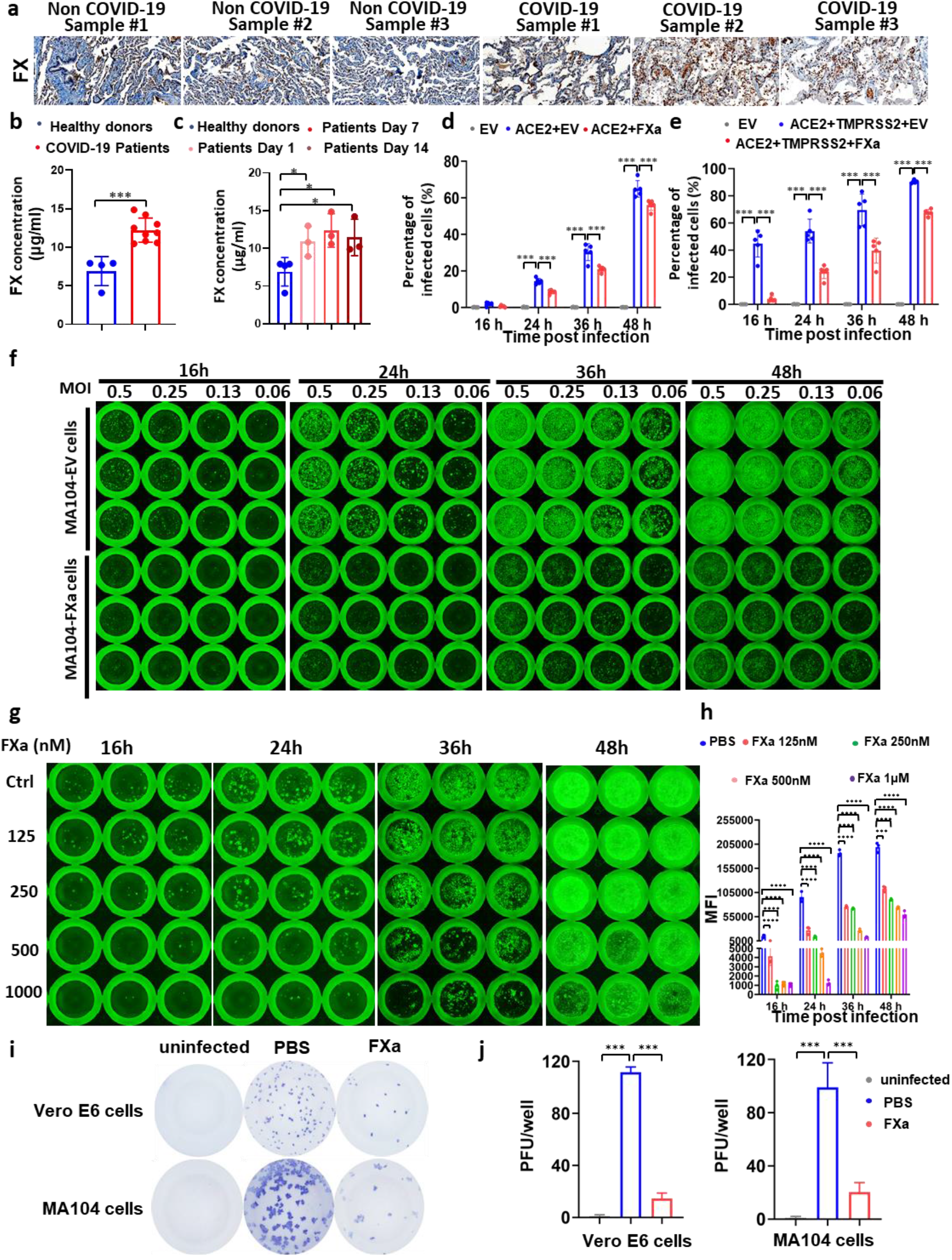
FXa inhibits wild-type SARS-CoV-2 infection by targeting viral particles. (**a and b**) FX protein levels in lungs (a) or serum (b) of COVID-19 patients vs. non-COVID-19 donors, using IHC (a) and ELISA (b), respectively. (**c**) FX concentrations in serum of COVID-19 patients as shown post diagnosis of infection were measured by ELISA. (**d and e**) HEK293T cells co-transfected with ACE2 and FXa or an EV in the absence (d) or presence (e) of TMPRSS2 were infected by VSV-SARS-CoV-2, and quantified by flow cytometry at 16, 24, 36, and 48 hpi. (**f**) MA104 cells transduced with FXa (MA104-FXa) or EV (MA104-EV) were infected by VSV-SARS-CoV-2 and imaged at 16, 24, 36, and 48 hpi by fluorescence microscopy. (**g and h**) VSV-SARS-CoV-2 was preincubated with FXa at indicated concentrations 1 hour before infection. Cells were imaged at 16, 24, 36, and 48 hpi by fluorescence microscopy (g) and the corresponding infectivity was measured by flow cytometry (h). (**i and j**). MA104 and Vero E6 cells were infected with live wild-type SARS-CoV-2. At 24 hpi, infectivity was measured by immune-plaque assay (i). (j) The summary data of (i). Experiments in i are representative of two independent experiments with similar data, and the other data are representative of at least three independent experiments. For all panels, error bars indicate standard deviation (SD), and statistical analyses were performed by Student’s t tests (b-e) and linear mixed model (h). *P ≤ 0.05; **P ≤ 0.01; ***P ≤ 0.001; ****P ≤ 0.0001; n.s, not significant.

To investigate the consequences of increased FXa during SARS-CoV-2 infection, we cloned FXa into the pCDH-mCherry vector and assessed its function using the vesicular stomatitis virus (VSV)-SARS-CoV-2 chimeric virus^18^. 293T cells were co-transfected with ACE2 and FXa or control empty vector (EV). After 24 hours, cells were infected by VSV-SARS-CoV-2. The percentage of infected cells (GFP positive cells) was examined at the indicated time points using flow cytometry. Surprisingly, at indicated hours post infection (hpi), compared to EV transfected group, the percentage of infected cells was decreased in FXa transfected group, indicating that the presence of FXa efficiently blocked viral infection (Fig. 1d). SARS-CoV-2 infection depends not only on ACE2 but also TMPRSS2^10^. When 293T cells were co-transfected with FXa, ACE2, and TMPRSS2, FXa again blocked viral infection, showing a more pronounced effect during the early time points in the presence of TMPRSS2 (Fig. 1e). To confirm our results, we generated an MA104 epithelial kidney cell line stably expressing FXa (MA104-FXa) (Extended Data Fig. 2a). The cells were infected by VSV-SARS-CoV-2 at different MOIs and the infectivity was determined at 16-48 hpi. We found that MA104-FXa cells showed markedly decreased infection at each time point and at all MOIs compared to MA104-EV control cells (Fig. 1f and Extended Data Fig. 2b). The viral titers of the supernatant from the infected MA104-FXa cells at 24 and 48 hpi were significantly decreased compared to that from MA104-EV cells, suggesting over-expressing FXa also impaired viral production (Extended Data Fig. 2c and d). We also compared the role of FXa with other SPs in viral infection of parental MA104 cells. Unlike pretreatment with furin, TMPRSS2, or trypsin, all of which increased VSV-SARS-CoV-2 infection, pre-treatment with FXa inhibited viral infection (Extended Data Fig. 3a and b). Consistent with this, the viral titer of the supernatant from FXa-treated MA104 cells infected with VSV-SARS-CoV-2 was significantly decreased, while the viral titer of infected cells treated with furin, TMPRSS2 or trypsin increased, compared to infected cells treated with vehicle control (PBS) (Extended Data Fig. 3c).

Furthermore, to determine if FXa blocks viral infection by targeting SARS-CoV-2 or host cells, we first constructed an FXa-Fc fusion protein expression plasmid and purified the protein from Chinese hamster ovary (CHO) cells. Then, we co-incubated VSV-SARS-CoV-2 with or without FXa-Fc fusion protein *in vitro* for 1 hour before adding the mixture into MA104 cells. The rate of infection was examined at the indicated timepoints. We found that pre-incubation of VSV-SARS-CoV-2 with FXa-Fc fusion protein significantly inhibited viral infection in a dose-dependent manner (from 62.5 pM to 1μM), suggesting that FXa could block viral infection by targeting SARS-CoV-2 (Fig. 1, g, h, and Extended Data Fig. 4). Of note, the inactivated FXa had no effect on viral infection (Extended Data Fig. 4). To determine the effects of FXa on viral production, we infected the MA104 cells with VSV-SARS-CoV-2 at a very low MOI (0.001) and with FXa protein at the indicated concentrations. We found that the viral titer showed a dose-dependent decrease with the increase of FXa protein, suggesting that FXa plays an essential role in inhibiting viral production (Extended Data Fig. 5a). To determine whether FXa also inhibits viral infection through interaction with host cells, we preincubated FXa-Fc fusion protein with MA104 cells for 1 hour and then washed out the medium before infecting cells with VSV-SARS-CoV-2. We found that FXa-Fc fusion protein pre-treatment with the MA104 cells did not significantly affect viral infection (Extended Data Fig. 5b and c). We next used live SARS-CoV-2 to infect Vero E6 and MA104-EV cells followed by quantitative assessment of viral load using an immuno-plaque assay. We found that pre-incubation FXa with live SARS-CoV-2 prior to infection significantly reduced viral infection in both Vero E6 and MA104 cells compared to buffer control, consistent with the above VSV-SARS-CoV-2 chimeric virus data (Fig. 1, i and j). The anti-viral effect of FXa against live SARS-CoV-2 infection was also confirmed with a traditional plaque assay with Vero E6 cells (Extended Data Fig. 5d). Together, our results showed that FXa, a serine protease that is upregulated following SARS-CoV-2 infection in host cells, inhibited viral infection and thus possessed an anti-viral activity, in distinct contrast to other serine proteases such as furin and TMPRSS2.

To study the mechanism(s) of the anti-viral activity of FXa, we compared the binding between FXa and various subunits of SARS-CoV-2 S protein. We found that FXa had the strongest binding affinity toward the full-length S protein and to a lesser extent to subunit S1, subunit S2 and the receptor binding domain (RBD) compared to the control Fc protein (Fig. 2a). Pull-down assay showed that FXa but not the Fc control protein co-precipitated with S protein (Fig. 2b). We then measured the binding affinity of FXa and the full-length S protein. The results showed that the binding affinity of FXa to S protein is in the nanogram range (Fig. 2c). These results suggest that FXa binds to the S protein, which might inhibit viral entry efficiency. Virus entry is followed by important conformational changes of viral proteins via cleavage of the S protein by host SPs. To determine whether FXa could cleave S protein, we incubated full-length S protein with FXa, followed by immunoblotting. Furin and TMPRSS2 served as positive controls, as they are known to induce functional conformational changes of S protein. We found that full-length S was cut into three fragments by FXa with the size of approximately 60 KD, then 50 KD and 29 KD (Fig. 2d), consistent with *in silico* prediction of two FXa cleavage sites on S protein, Ile-(Asp/Glu)-Gly-Arg (R1000) and Gly-Arg (R567) (Fig. 2e). This cleavage pattern was in contrast to that of furin and TMPRSS2, which both cut full-length S protein into the ~80 KD subunit S1. Cleavage by FXa did not produce the ~80 KD subunit S1.

**Fig. 2.**
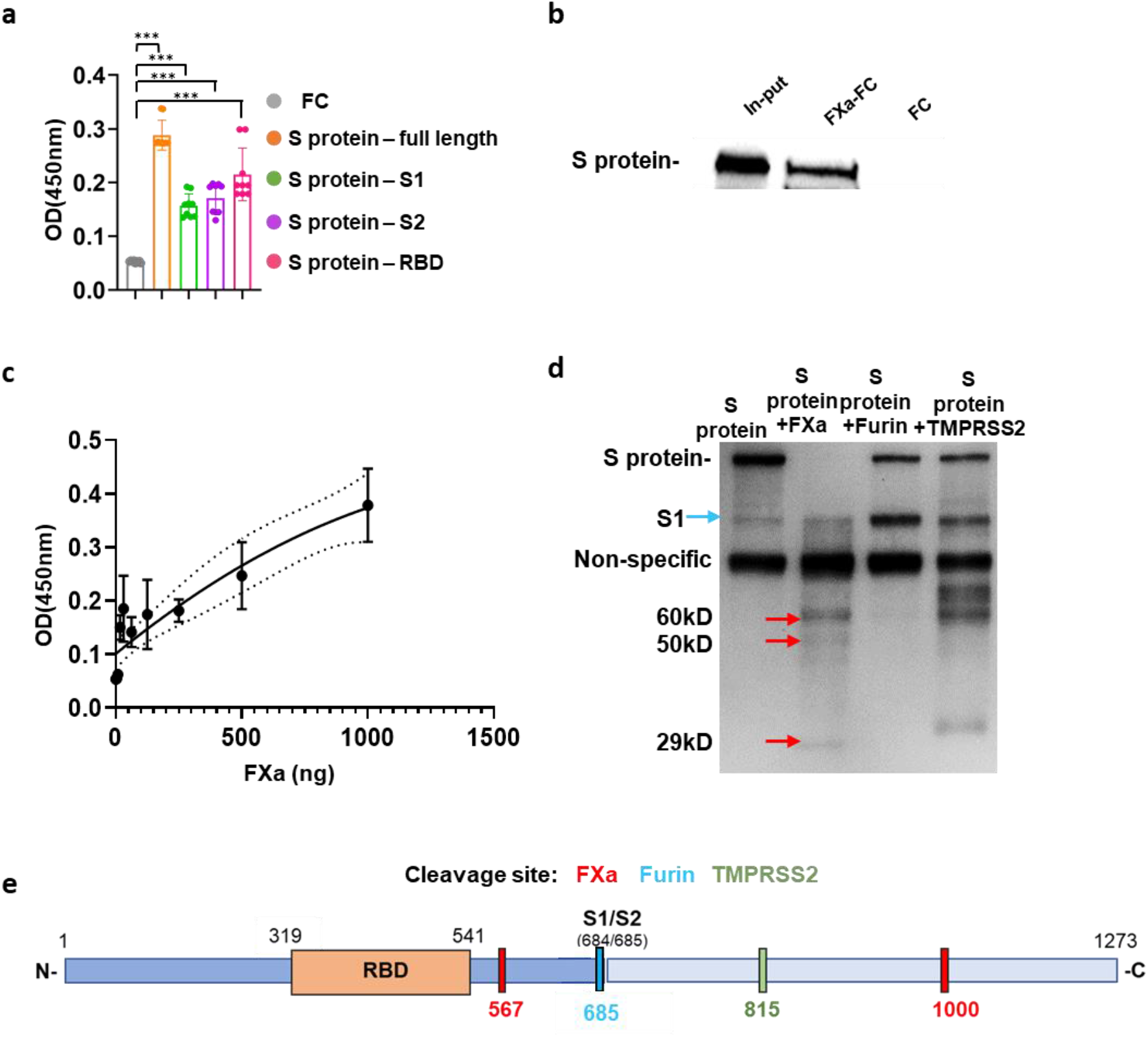
FXa suppresses viral entry by binding to and cleaving the SARS-CoV-2 S protein. (**a**) The binding affinity of FXa with full-length wild-type S protein, subunit S1, subunit S2, and RBD was quantified by ELISA. (**b and c**) The interaction between FXa protein and full-length S protein was examined by pull-down assay (b). The binding affinity at indicated concentrations of FXa was measured by ELISA (c). (**d**) The cleavage of S protein by furin, TMPRSS2, and FXa was analyzed by immunoblotting. (**e**). Schema of the cleavage sites for furin, TMPRSS2 and FXa on the full-length S protein. (**f**) 293 T cells were co-transfected with a S protein expression plasmid and various amounts of an FXa expression plasmid. S protein cleavage by FXa inside of cells was analyzed by immunoblotting. All data are representative of at least three independent experiments. Experiments in b and d are representative of three independent experiments with similar data. For all panels, error bars indicate SD, and statistical analyses were performed by one-way ANOVA models. ***P ≤ 0.001.

Given that FXa could cut S protein into different sizes, we used ELISA to determine whether cleavage by FXa affected binding of the S protein to ACE2. The ELISA data indicated that S protein pre-treated with FXa resulted in decreased binding affinity to ACE2 (Fig. 3a). Flow cytometry further confirmed this result, demonstrate that FXa-pre-treated S protein could not efficiently bind to ACE2-expressing HEK293T cells (Fig. 3, b and c). We also showed that FXa still bound to S protein when S protein was already bound to ACE2 (Fig. 3, d, e and f), yet FXa still cleaved the S protein bound to ACE2 (Extended Data Fig. 6). Overall, our results indicate that FXa could be an efficient inhibitor of viral entry via its interaction with S protein.

**Fig. 3.**
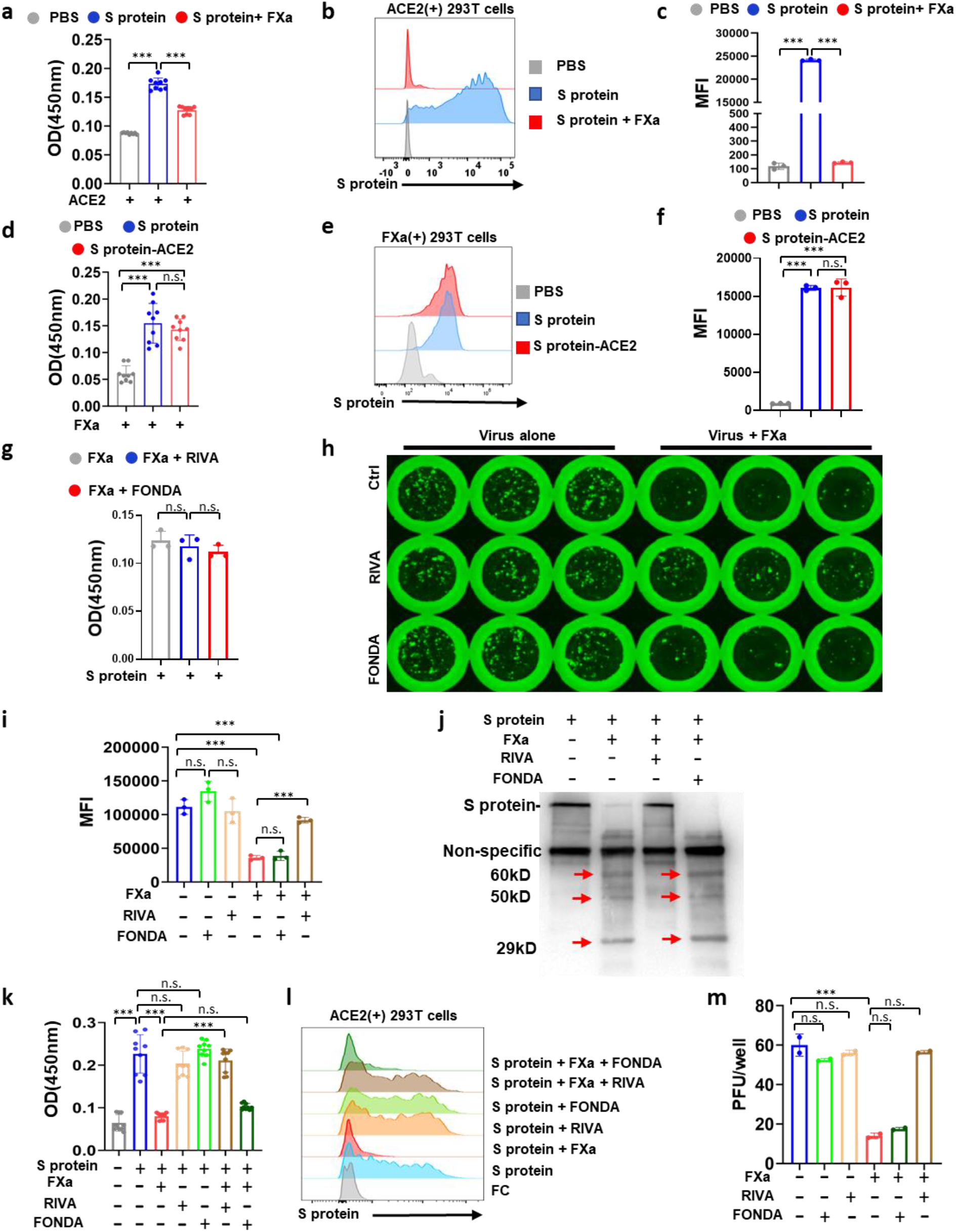
FXa cleavage blocks the binding between S protein and ACE2. (**a**) The binding between ACE2 with S protein or FXa-pretreated S protein was measured by ELISA. (**b and c**) The binding between S protein or FXa-pretreated S protein and ACE2 expressed on 293T cells was measured by flow cytometry (b and c, summary data). (**d**) The binding of FXa with S protein, S protein-ACE2 complex, or PBS control, assessed by ELISA. (**e** and **f**) The binding between the S protein or the S protein-ACE2 complex with FXa expressed on 293T cells. PBS served as control for S protein and S protein-ACE2 complex. (**g**) The effect of RIVA or FONDA on the binding of FXa with S protein was measured by ELISA. (**h and i**) The infectivity of FXa-pretreated vs. untreated VSV-SARS-CoV-2 in MA104 cells in the presence or absence of RIVA or FONDA was examined with fluorescent microscopy (h) and flow cytometry (i). (**j**) S protein cleavage by FXa in the presence or absence of RIVA or FONDA was examined by immunoblot. (**k and l**) FXa pretreated with or without RIVA or FONDA was incubated with S protein, followed by assessing the binding capability of these S proteins with ACE2 coated on a plate (j) or expressed on 293T cells (l). (**m**) The infectivity of FXa-pretreated vs. untreated live SARS-CoV-2 in MA104 cells in the presence or absence of RIVA or FONDA was examined using an immune-plaque assay. All data are representative of at least three independent experiments. Experiments in h and j are representative of three independent experiments with similar data. For all panels, error bars indicate SD, and statistical analyses were performed by one-way ANOVA models. ***P ≤ 0.001; n.s, not significant.

An emergent SARS-CoV-2 strain has been found to substitute aspartic acid–614 for glycine (D614G) in the S protein^19^. Thus, we tested whether FXa had similar functional interactions with the D614G S protein. FXa could still bind with and cleave the D614G S protein (Extended Data Fig. 7a-c), and the binding affinity between ACE2 and D614G S protein decreased if S protein was pretreated with FXa (Extended Data Fig. 7d and e).

COVID-19 patients with an increased risk of thrombosis are treated with direct FXa inhibitors (e.g., rivaroxaban) or indirect inhibitors (e.g., fondaparinux)^14,20^. We therefore asked if rivaroxaban (RIVA) or fondaparinux (FONDA) affect the anti-viral activity of FXa. Neither RIVA nor FONDA blocked the binding of FXa to S protein (Fig. 3g), and neither drug alone had any effect on VSV-SARS-CoV-2 infectivity; however, the direct FXa inhibitor RIVA blocked FXa-induced anti-viral activity, whereas the indirect FXa inhibitor FONDA did not (Fig. 3, h and i). We measured viral titers in the supernatants collected from this inhibitor experiment, which confirmed that RIVA but not FONDA significantly reduced FXa-induced anti-viral activity (Extended Data Fig. 8a). Furthermore, cleavage assay showed that the direct FXa inhibitor RIVA but not the indirect inhibitor FONDA inhibited cleavage of S protein by FXa (Fig. 3j). Consistent with this, pretreatment of the mixture of S protein and FXa with RIVA or FONDA, followed by incubation with ACE2, showed that RIVA but not FONDA significantly diminished the effect of FXa on inhibiting the binding of S protein to ACE2 as demonstrated by ELISA, presumably by inhibiting cleavage of S protein by RIVA but not by FONDA (Fig. 3k). The ELISA results in Figure 3k were validated by flow cytometry analysis (Fig. 3l and Extended Data Fig. 8b). Our data indicate that the direct rather than indirect inhibitor of FXa could diminish FXa-mediated blockade of viral entry by inhibiting the cleavage of S protein by FXa so that intact S protein can efficiently bind to ACE2. We repeated these experiments with live SARS-CoV-2 and MA104 cells demonstrating identical results (Fig. 3m).

To evaluate the potential effect of FXa *in vivo*, we used humanized K18-hACE2 mice as an infection model of SARS-CoV-2^21,22^ and inoculated them with 3×10^5^ pfu SARS-CoV-2, followed by intranasal administration of the FXa-Fc protein or two controls, saline and the Fc protein. The body weight of the mice was monitored. We found that the majority of mice in the untreated and Fc-treated groups exhibited a dramatic decrease in their body weight at day 5 and were euthanized at that time or shortly thereafter, while three of the five mice treated with FXa-Fc started the recovery of their body weight at day 7 (Fig. 4a). The FXa-Fc treated group lived significantly longer than the two control groups, with no difference between the control groups (Fig. 4b). We isolated RNA and used quantitative real-time PCR to measure the viral load in trachea, lung, and brain tissues. Viral load in the FXa-Fc treated group was approximately 1,000-fold lower than that of the two control groups, indicating that FXa-Fc significantly restricted SARS-CoV-2 infection *in vivo* (Fig. 4, c, d, and e). Consistent with this, IHC showed that expression of viral nucleocapsid protein (NP) was also markedly decreased in the brain and lung tissues from the FXa-Fc treated group compared to the untreated and Fc-treated group (Fig. 4f). The histological study showed that FXa-Fc-treated mice had more intact lung structure and less pathological damage compared to the two control groups (Fig. 4f). To evaluate the effects of the two types of FXa inhibitors, the direct inhibitor RIVA and the indirect inhibitor FONDA *in vivo*, we were administrated into FXa-treated (intranasally) SARS-CoV-2-infected mice. Consistent with the *in vitro* data, we found that direct FXa inhibitor RIVA significantly blocked the anti-viral and survival advantage afforded by intranasal administration of FXa-Fc, while the indirect FXa inhibitor FONDA had no significant effect on the anti-viral and survival advantage afforded by intranasal administration of FXa-Fc alone (Fig. 5, a-e). These *in vivo* results with live SARS-CoV-2 provide preclinical support for the use of an indirect FXa inhibitor such as FONDA as an anti-coagulant when preventing or treating thrombotic complications of COVID-19, while avoiding the use of a direct FXa inhibitor such as the anti-coagulant RIVA under similar clinical circumstances.

**Fig. 4.**
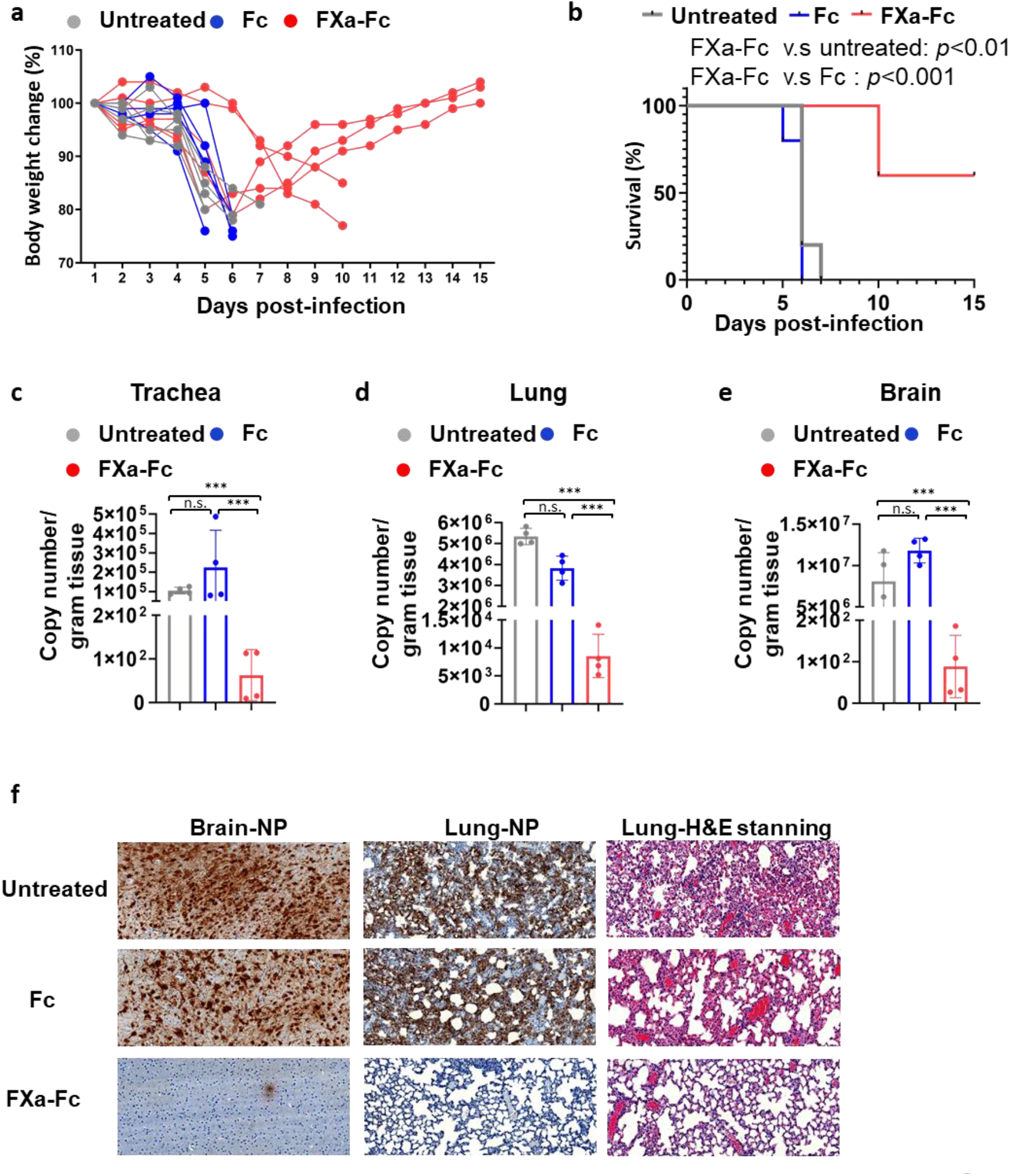
The effect of FXa protein on WT SARS-CoV-2 infection in a K18-hACE2 mouse model of COVID-19. (**a and b**) Body weight (a) and survival (b) of mice infected with 3×10^5^ PFU SARS-CoV-2 WTstrain and treated with or without FXa-Fc fusion protein. Fc-protein was used as control. (**c-e**) Viral load in the tracheas (c), lungs (d), and brains (e) of mice treated with or without FXa-Fc fusion protein or Fc control was assessed by Q-PCR. (**f**) The presence of SARS-CoV-2 was determined using IHC staining with an antibody against viral nucleocapsid protein (NP) in the brain and lung of mice treated with FXa-Fc or Fc-protein. Pathological analysis of the lung of mice treated with or without FXa-Fc or Fc-protein as performed by H&E staining. For all panels, error bars indicate SD, and statistical analyses were performed by one-way ANOVA models (c, d, e) and log-rank test (b). ***P ≤ 0.001; n.s, not significant.

**Fig. 5.**
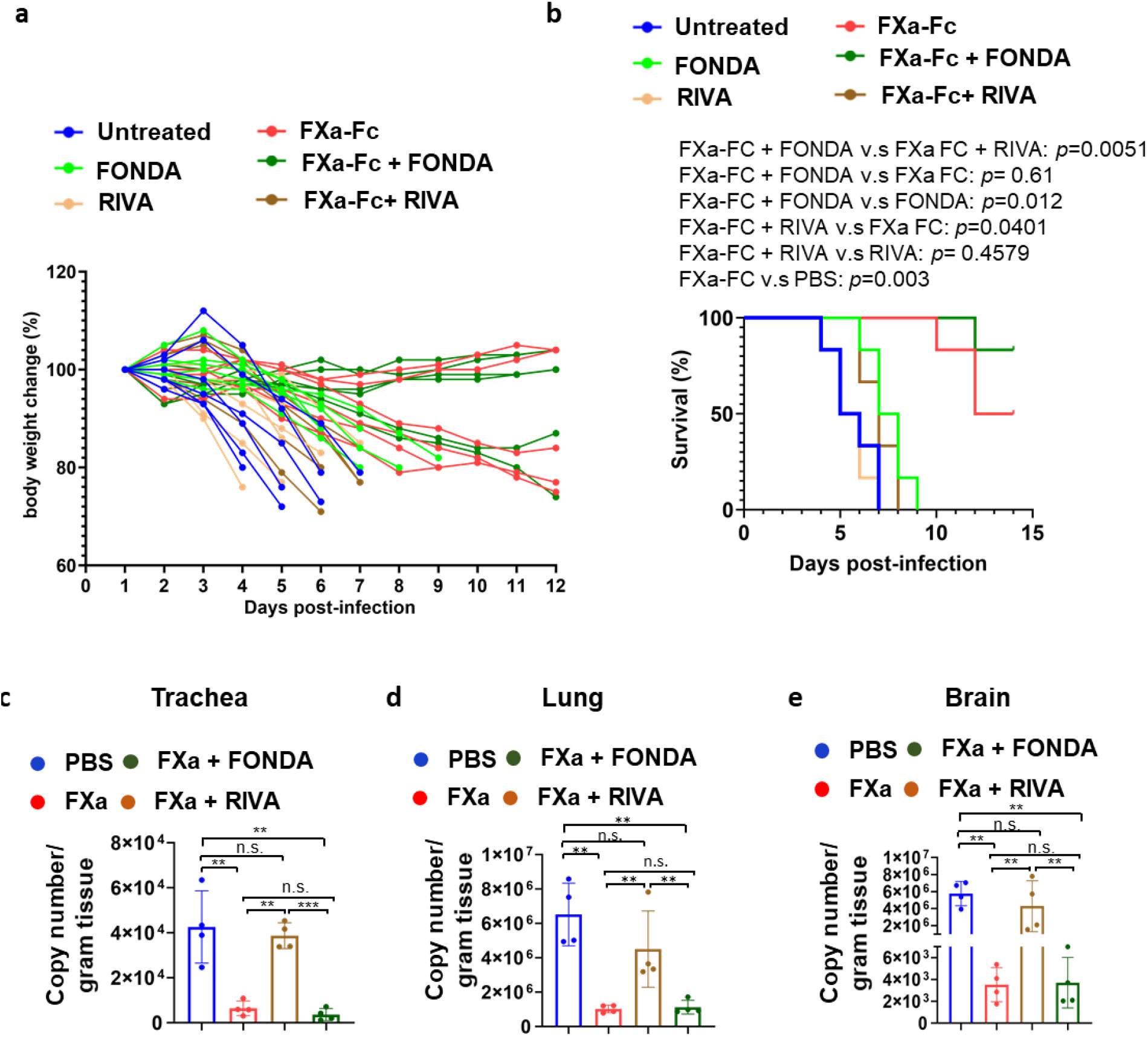
The effect of the direct FXa inhibitor RIVA and the indirect inhibitor FONDA on FXa-mediated protection of K18-hACE2 mice from WT SARS-CoV-2 infection. (**a and b**) Body weight (a) and survival (b) of mice infected with 3×10^5^ PFU SARS-CoV-2 (WT-1) and treated with or without FXa-Fc in the presence or absence of RIVA or FONDA. (**c-e**) Viral load in the trachea (c), lung (d), and brain (e) of mice treated with or without FXa-Fc in the presence or absence of RIVA or FONDA was assessed by Q-PCR. For all panels, error bars indicate SD, and statistical analyses were performed by one-way ANOVA models (c, d, e) and log-rank test (b). **P ≤ 0.01; ***P ≤ 0.001; n.s, not significant.

A variant of SARS-CoV-2 (known as B.1.1.7), which was emerged with a large number of mutations, may be associated with an increased transmissibility and risk of death compared with other variants^23,24^. By analyzing the S protein sequence of the B.1.1.7 variant, we found there is a nonsynonymous mutation A570D in the S protein close to the predicted FXa cleavage site. In order to evaluate the effect of the anti-viral function of FXa on the variant, we infected the Vero E6 and MA104 cells with the original emergent SARS-CoV-2 (wild-type; WT) or B.1.1.7 virus pretreated with different concentrations of FXa. The immuno-plaque results showed that the anti-viral effect of FXa was significantly and dramatically decreased against B.1.1.7 variant infection compared to the WT infection (Fig. 6a). Furthermore, the results were confirmed by viral infection with various MOIs, which showed that FXa could still block WT infection even at a very high MOI (MOI=8), but had little effect on B.1.1.7 infection blockade at the same MOI (Fig. 6b). To figure out the mechanism of this difference, we detected the binding affinity between FXa and WT S protein or B.1.1.7 S protein. Both ELISA and flow cytometry results showed that FXa had a significantly lower binding affinity with B.1.1.7 S protein compared to that with WT S protein (Fig. 6, c and d), indicating that the mutants on B.1.1.7 S protein blocked FXa anti-viral effect by affecting the binding affinity of FXa to the S protein. We also tested the effects of the direct and indirect FXa inhibitors in the B.1.1.7 variant infection using the WT strain as control. We found that RIVA could efficiently block the anti-viral effect of FXa in a dose-dependent manner starting as low as 0.05 μg/ml dose, while FONDA did not even at the high dose of 50 μg/ml, against the B.1.1.7 variant infection in both Vero E6 and MA104 hosts (Extended Data Fig. 9a and b). These data were validated in similar experiments with gradient concentrations of FXa in both Vero E6 and MA104 hosts (Extended Data Fig. 10 and 11).

**Fig. 6.**
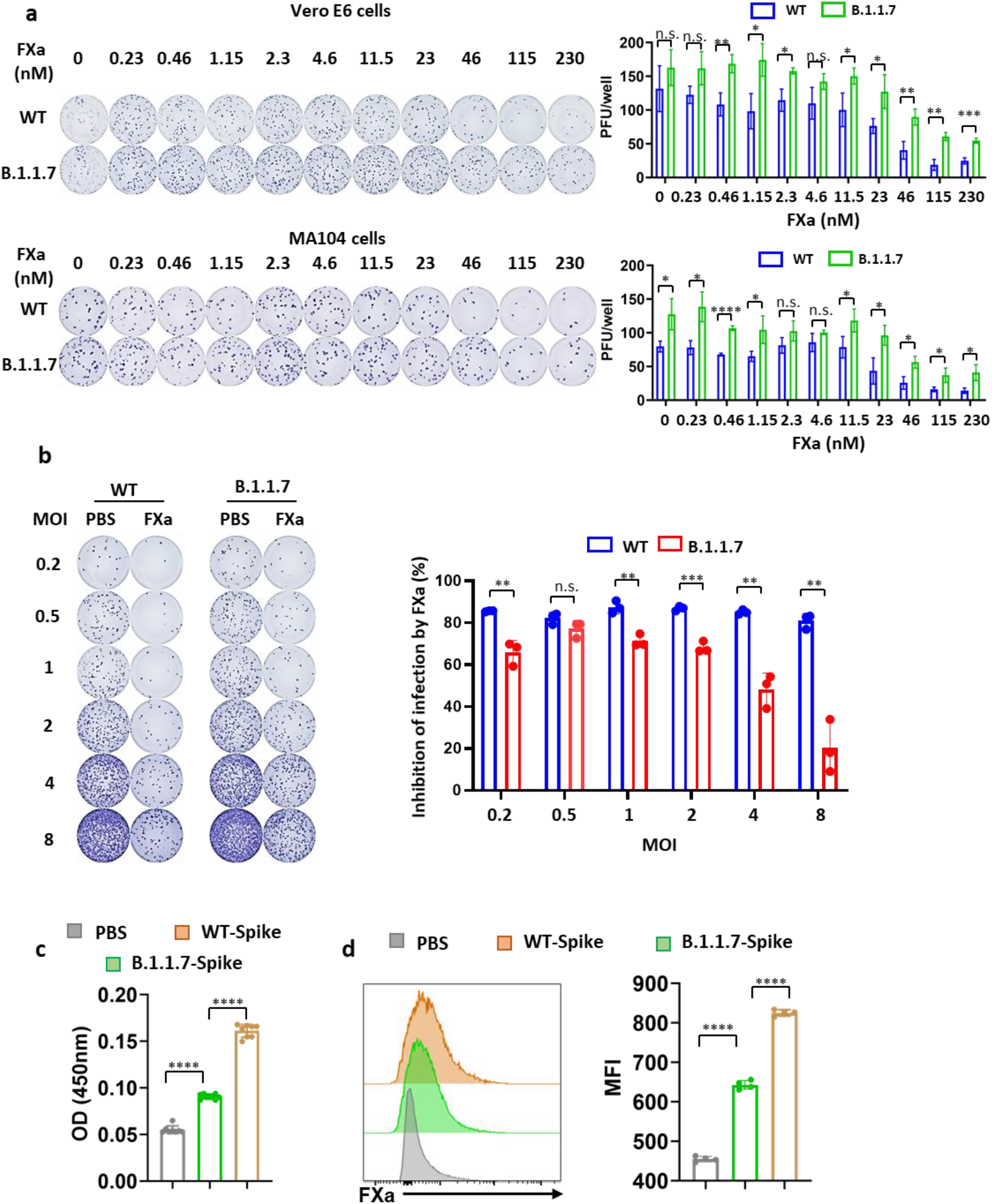
FXa blocks the B.1.1.7 variant infection less efficiently compared to the WT SARS-CoV-2 due to different binding affinity to the corresponding S protein. (**a**) MA104 and Vero E6 cells were infected with either the live wild-type or the B.1.1.7 variant of SARS-CoV-2 pretreated with different doses of FXa. At 24 hpi, infectivity was measured by immune-plaque assay. The representative infection and the summary data are presented at the left and right, respectively. **(b)** MA104 and Vero E6 cells were infected with live wild-type or the B.1.1.7 SARS-CoV-2 variant pretreated with FXa at different MOIs. At 24 hpi, infectivity was measured by immune-plaque assay (left panel) and the infection inhibition ratio induced by FXa at different MOIs are summarized (right panel). **(c)** The binding FXa with wild-type S protein or B.1.1.7 variant S protein, assessed by ELISA. (**d**) The binding between wild-type S protein or B.1.1.7 variant S protein with FXa expressed on 293T cells, assessed by flow cytometry. PBS serves as control. For all panels, error bars indicate SD, and statistical analyses were performed by one-way ANOVA models. *P ≤ 0.05; **P ≤ 0.01; ***P ≤ 0.001; ****P ≤ 0.0001; n.s, not significant.

Next, we compared the anti-viral effect of FXa against WT and B.1.1.7 infection *in vivo*. Consistent with the *in vitro* data, we found that the anti-viral and survival advantage afforded by FXa-Fc was abolished or significantly decreased in the B.1.1.7 variant-infected group compared to the WT-infected group (Fig. 7, a-d). Collectively, our data demonstrate that the variant B.1.1.7 strain with a mutated spike protein was resistant to the FXa inhibition *in vitro* and *in vivo* compared to the WT strain, which might explain the higher transmission^25–27^ and mortality^28^ rates for this dangerous variant of concern.

**Fig. 7.**
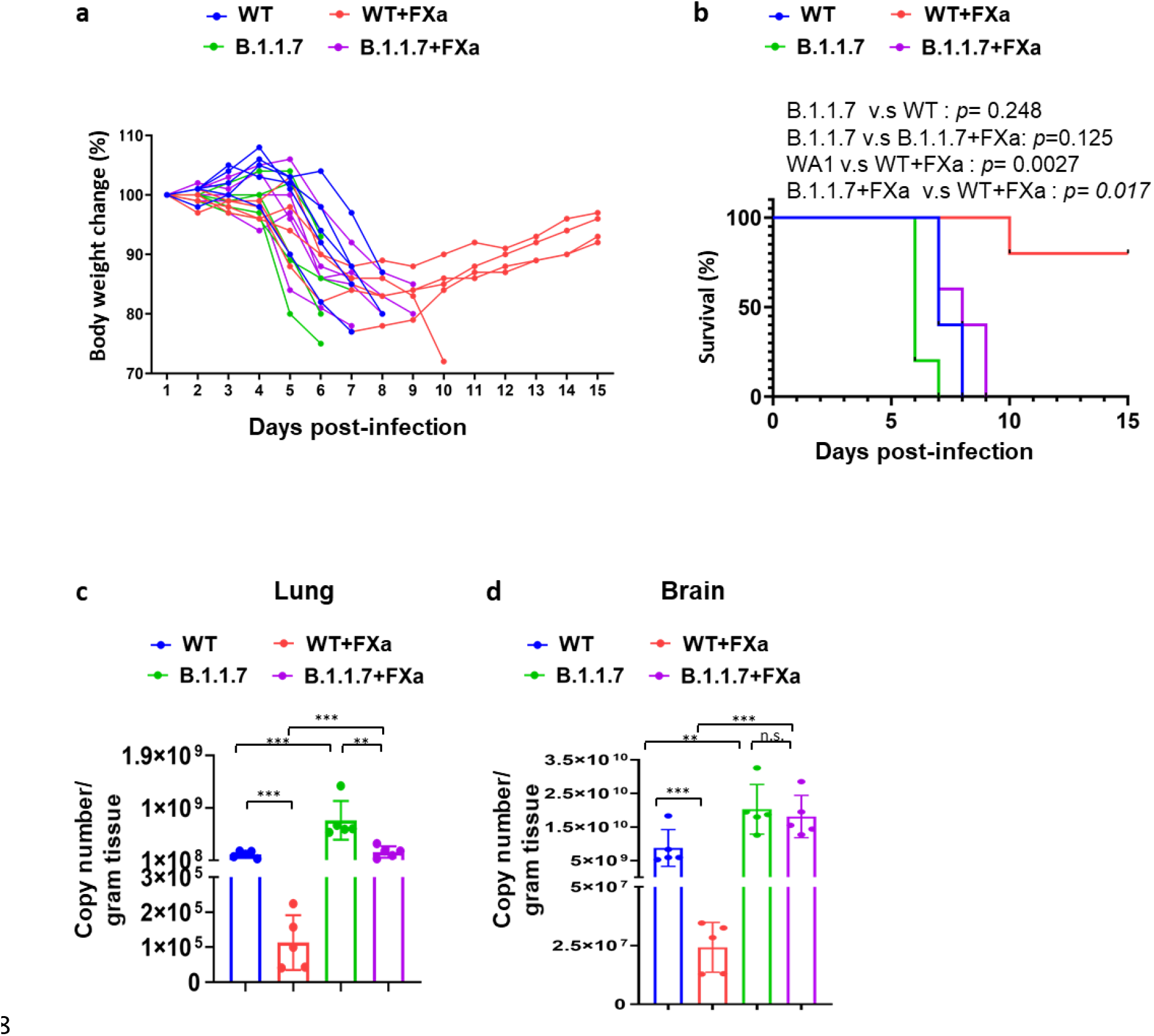
The effect of FXa protein on live B.1.1.7 variant infection in in a lethal humanized hACE2 mouse model of SARS-CoV-2 infection. (**a and b**) Body weight (a) and survival (b) of mice infected with 3×10^5^ PFU wild-type SARS-CoV-2 or B.1.1.7 variant and treated with or without FXa-Fc fusion protein. (**c and d**) Viral load in the lung (c) and brain (d) of mice treated with or without FXa-Fc fusion protein was assessed by quantitative PCR. For all panels, error bars indicate SD, and statistical analyses were performed by one-way ANOVA models (c and d) and log-rank test (b). ***P ≤ 0.001; n.s, not significant.

## Discussion

Here we identify a novel mechanism of human host anti-viral defense involving FXa at the time of SARS-CoV-2 infection, which binds to and cleaves viral S protein, blocking viral entry into host cells. Exogenous administration of FXa reduced viral load and protected a humanized hACE2 mouse model of COVID-19 from lethal infection, an effect that was attenuated by a direct but not indirect FXa inhibitor and anti-coagulant, which has implications for clinical therapeutic responses.

SARS-CoV-2 is a newly emergent human pathogen that utilizes the ACE2 receptor to enter host cells. Proteolytic processing S protein by SPs such TMPRSS2, furin, and trypsin enhances the binding affinity between host ACE2 and the processed S protein^10^. In the course of infection, unlike the immune tolerance exhibited by bats, in human SARS-CoV-2 at times excessively activates the inflammatory components of the immune system of humans leading to the cytokine release syndrome ^29^, which can be fatal in some, yet non-existent in others with the same exposure ^30^. Therefore, identification of the body’s natural defense mechanisms against SARS-CoV-2 is important for developing effective prevention as well as therapeutic strategies. Our report here of a natural defense mechanism involving the binding to and cleavage of the SARS-CoV-2 S protein by FXa, effectively blocking viral entry into host cells, may lead to the development of such strategies. Like TMPRSS2 and furin, FXa belongs to the family of SPs that each cleave S protein, but have different cleavage targets and subsequent effects on viral entry into host cells. TMPRSS2 and furin cleave the S protein at the S1/S2 site of at RRAR (685R), followed by the S2’ site KPSKR (R856), resulting in priming the SARS-CoV-2 fusion step. How the FXa cleavage of the S protein blocks entry of SARS-CoV-2 into host cells is currently unknown. But our *in silico* modeling, which is supported by experimental evidence, indicates that the cleavage sites of FXa on S protein are at Ile-(Asp/Glu)-Gly-Arg (R1000) or Gly-Arg (R567), distinct from the S1/S2 or the S2’ site. This likely results in a unique conformational change of the S protein when forming a syncytium^31^. The most conserved region of the RBD is between amino acids 306 to 527, which is close to the FXa cleavage site (R567)^8^. As such, the cleavage by FXa near the RBD may adversely impact the conserved conformation of the RBD. The S protein is a type 1 viral fusion protein with two conserved heptad repeat regions, HR-N (916-950) and HR-C (1150-1185), which may form a 6-helix bundle allowing for a better fusion between the viral and host cell membranes^32,33^. The two likely cleavage sites of FXa are located within the HR-N and the HR-C repeat and could impair the formation of a hetero-stranded complex, impairing the fusion process.

The B.1.1.7 variant of concern emerged in the United Kingdom in September 2020 and has demonstrated higher transmissibility^25–27^ and mortality^28^. It has been shown that some vaccines and neutralizing antibodies have lower efficacy from B.1.1.7 variant infection than WT strain infection^34–38^. However, the reason for its poor outcomes remained elusive^39^. Our description of the previously unknown FXa barrier to SARS-COV-2 infection and its differential effect on the B.1.1.7 variant is a possible, or at least partial, explanation. Our data showed that the variant B.1.1.7 strain with a mutated spike protein was resistant to the FXa inhibition *in vitro* and *in vivo* compared to the WT strain. Therefore, the host anti-viral defense system depending on FXa maybe not strong enough to protect people from B.1.1.7 variant infection. The basis for the variant differential could be the A570D mutation, which only emerged as a dominant change in the B.1.1.7 variant near the FXa cleavage site. FXa showed a significantly lower binding affinity with B.1.1.7 S protein compared to that with WT S protein, which might be affected by the A570D mutation in the B.1.1.7 S protein. But the spike protein structure is complex and dynamic, which means more distal differences could also play a role.

FXa is required for the conversion of prothrombin to thrombin in the clotting cascade^12^, and may have a role in inflammation^40^. Both of these processes are dysregulated in some patients with COVID-19^30,41^. Although it is likely that the viral defense mechanism that we have described here is active and important in controlling SARS-CoV-2 infection in asymptomatic and mildly symptomatic individuals, the endogenous overexpression of FXa during serious SARS-CoV-2 infection may also contribute to the pathogenesis and complications of COVID-19, especially the thrombotic events^41,42^. Therefore, the use of FXa as a therapeutic agent is straightforward. There could be at least two alternative pathways forward in considering FXa as a therapeutic agent for severe SARS-CoV-2 infection. The first would be to co-administer an indirect FXa inhibitor as an anti-coagulant in combination with the FXa-Fc fusion protein described in this report or with a recombinant FXa, because we show that the indirect FXa inhibitor fondaparinux does not interfere with FXa’s cleavage of S protein or its therapeutic effect against the live virus *in vivo*. The second approach would be to modify the chemical structure of FXa such that its enzymatic activity for cleavage of the S protein is retained while that of prothrombin conversion is lost. This latter approach might also be considered as a preventative approach for individuals who are not vaccinated and are highly susceptible to severe COVID-19 or those same individuals who are vaccinated yet fail to develop effective immunity to SARS-CoV-2.

There are four clinically approved direct FXa inhibitors, including rivaroxaban, apixaban, edoxaban as well as betrixaban) and one indirect FXa inhibitor fondaparinux for use as anti-thrombotic agents in patients with hypercoagulable states^43^. Our study showed that in the presence of a direct inhibitor of FXa, rivaroxaban, the anti-SARS-CoV-2 activity of FXa is affected not by the binding of the S protein but by the cleavage of the S protein. Further, rivaroxaban completely abrogates the decrease in viral load and the protective effect against lethal SARS-CoV-2 infection conferred by exogenous FXa. Our study suggests that the use of direct FXa inhibitors such as rivaroxaban, in patients highly susceptible for severe COVID-19 as is currently being evaluated in a number of studies^14^, should likely proceed with caution as it is at least conceivable from our work that such studies could result in an increase in viral load. Importantly, the FXa indirect inhibitor, fondaparinux, which we found to not block the protective effects of FXa against SARS-CoV-2, has been found to be safe and efficacious for venous thrombosis prophylaxis in hospitalized COVID-19 patients^20^.

The factors responsible for predicting clinical outcomes in patients stricken with COVID-19 remain incomplete^44^. Here we show that the precursor FX of FXa is upregulated in COVID-19 patients. We identify a new mechanism of anti-viral defense in human and prove its importance in a transgenic animal model that mimics the human disease. Our work would suggest that future studies should examine the quality and quantity of FXa enzymatic activity against the SARS-CoV-2 S protein to determine if it can improve our understanding of who may be most susceptible to SARS-CoV-2 infection and who might be treated with an FXa indirect anti-coagulant inhibitor.

## Methods

### Patient sample collection

Patients were collected and tested positive for SARS-CoV-2 at City of Hope. Autopsy samples were provided by Dr. Ross Zumwalt at the University of New Mexico School of Medicine. The concentration of FXa in the serum of patient samples was measured using ELISA (LS-F10420-1, LSBIO). The protocols for human specimen collection were approved by the institutional review board of City of Hope.

### Cells

Monkey kidney epithelial-derived MA104 cells were maintained in medium 199 supplemented with 10% FBS, penicillin (100 U/ml), and streptomycin (100 μg/ml). To overexpress FXa in MA104 cells, the cells were infected with lentivirus encoding FXa to generate MA104-FXa cells. Monkey kidney epithelium-derived Vero cells, Vero E6 cells, human embryonic kidney-derived HEK293T cells, and Chinese hamster ovary (CHO) cells were cultured in DMEM with 10% FBS, penicillin (100 U/ml), and streptomycin (100 μg/ml). All cell lines were routinely tested to confirm absence of mycoplasma using the MycoAlert Plus Mycoplasma Detection Kit from Lonza (Walkersville, MD).

### VSV-SARS-CoV-2 infection

The VSV-SARS-CoV-2 chimeric virus expressing GFP was kindly provided by Sean Whelan at Washington University School of Medicine. The virus is decorated with SARS-CoV-2 S protein in place of the native glycoprotein (G) ^1^. For VSV-SARS-CoV-2 infection, MA104 cells were seeded 24 hours before the infection at a confluency of 70% in a 96-well plate. VSV-SARS-CoV-2 virus and varying amounts (12.5 μg/ml, 25 μg/ml, 50 μg/ml, and 100 μg/ml) of the FXa-Fc fusion protein were co-incubated at 37°C for 1 hour and then were added to the cells. To assess the effect of FXa inhibitors, FXa protein was preincubated with or without 50 μg/ml rivaroxaban or fondaparinux separately for 1 hour at room temperature. Infectivity was assessed by detecting GFP fluorescence using a Zeiss fluorescence microscope (AXIO observer 7) and/or determined by the percentage of GFP(+) cells analyzed using a Fortessa X20 flow cytometer (BD Biosciences) at 16, 24, 36, and 48 hours post infection (hpi). To determine viral production, Vero cells were pre-seeded for 24 hours and infected with the supernatants collected from MA104 cells infected by VSV-SARS-CoV at 24 or 48 hpi. The supernatants were diluted by 5-fold before the viral production assay.

### Generation and purification of FXa

CHO cells were transduced with a pCDH lentiviral vector expressing FXa to produce the FXa-Fc fusion protein for functionality assays. For this purpose, FXa fused with human IgG4 was reconstructed as previously reported ^2^. mCherry was co-expressed with FXa for FACS-sorting to purify transduced cells using a FACS Aria II cell sorter (BD Biosciences, San Jose, CA, USA). Conditional supernatants from lentivirus-infected CHO cells sorted by FACS were used to purify the FXa-Fc fusion protein using a protein G column (89927, Thermo Fisher). For *in vivo* testing, the protein G column-purified FXa-Fc fusion protein was desalted using fast protein liquid chromatography (FPLC).

### SARS-CoV-2 neutralization, cell infection, plaque assay, and immunoplaque assay

The following reagent was obtained through BEI Resources, NIAID, NIH: SARS-Related Coronavirus 2, Isolate USA-WA1/2020, NR-52281 (wild-type, WT) and SARS-Related Coronavirus 2, Isolate USA/CA_CDC_5574/2020, NR-54011 (B.1.1.7). Virus isolates were passaged in Vero E6 cells (ATCC CRL-1586) as previously described^3^. Virus concentration was determined using immunoplaque assay (also called focus forming assay) ^4^. For the plaque assay, 120 pfu SARS-CoV-2 was incubated with diluted sera for 2 hours at 37 °C. Then Vero E6 cells were infected with 250 μl virus-sera mixture for 1 hour. After infection, the medium containing virus was removed, and overlay medium containing FBS-free DMEM and 2% low-melting point agarose was added. At 72 hours post infection, infected cells were fixed by 4% paraformaldehyde (PFA) overnight, and stained with 0.2% crystal violet. For the immunoplaque assay, 100 pfu of live SARS-CoV-2 variants were incubated with diluted sera for 1 hour and then the virus antibody mixture was added to Vero E6 cells for 1 hour at 37°C. After 1 hour the virus containing medium was removed, overlayed with medium containing methylcellulose and 2% FBS DMEM, and incubated at 37°C. At 24 hours after infection, infected cells were fixed by 4% paraformaldehyde for 20 minutes at room temperature and then permeabilized by 0.5% Triton X-100/ PBS solution for 210 minutes at room temperature. SARS-CoV-2 viral nucleocapsid protein (NP) was detected using the anti-NP protein antibody (PA5-81794, Thermo Fisher) diluted 1:10000 in 0.1% tween-20/1%BSA/PBS solution as a primary antibody, followed by detecting with an anti-rabbit secondary antibody (ab6721, Abcam) at a 1:20,000 dilution. Plates were washed three times between antibody solutions using 0.5% tween-20 in PBS. The plates were developed using TrueBlue Peroxidase Substrate (5510-0030, Sera Care) and then scanned using Immunospot S6 Sentry (C.T.L Analyzers). Neutralization titers for the immunoplaque assay are defined as a 50% reduction in plaque forming units relative to the untreated wells.

### Assessment of binding between S protein and FXa using ELISA

Full-length coronavirus S protein with His tag (500 ng) (40589-V08B1, Sino Biological), coronavirus S protein S1 subunit with His tag (500 ng) (40591-V08B1, Sino Biological), coronavirus S protein S2 subunit with His tag (500 ng) (40070-V08B, Sino Biological), and coronavirus S protein RBD with His tag (500 ng) (40592-V08B-B, Sino Biological) were used as coating reagents in 96-well plates (3361, Corning). Coated plates were incubated with FXa protein (1 μg/ml) for 2 hours at room temperature. FXa-HRP conjugated anti-human Fc antibody (05-4220, Invitrogen) was used as a detecting antibody. Absorbance was measured at OD450 nm using a Multiskan™ FC Microplate Photometer (Fisher Scientific).

### Pull-down assay

HEK293T cells were transduced with a pCDH lentiviral vector expressing the full-length spike (S) protein for 48 hours. The cells were lysed and incubated with FXa-Fc or Fc (10 μg/ml) for 3 hours, then 20 μl protein A agarose resin beads (P-400-25, Invitrogen) were added and incubated overnight. After incubation, the beads were washed and collected. Protein binding between FXa-Fc or Fc and S protein was detected by immunoblotting using an anti-S protein antibody (ab272504, Abcam).

### Cleavage assay

One microgram of full-length S protein was treated with 1 μg of FXa (P8010L, NEB), furin (P8077S, NEB) or TMPRSS2 (TMPRSS2-1856H, Creative BioMart) protein for 3 hours following the manufacturers’ instructions. Cleavage was detected using immunoblotting with an anti-S protein antibody. For inhibitor assays, FXa protein was preincubated with or without 50 μg/ml rivaroxaban or fondaparinux separately for 1 hour in advance at room temperature, then treated with SPs and detected as above. For cleavage assays using S protein-ACE2 complex, 1 μg S protein and 1 μg ACE2 were pre-incubated 1 hour prior to incubation with FXa, then treated with SPs and detected as above. Of note, we used different buffer conditions for all binding assays and cleavage assays.

### Assessment of binding between S protein and FXa using flow cytometry

HEK293T cells were transduced with lentiviral vector expressing FXa for 48 hours. The cells were incubated with 10 μg/ml full-length S protein for 20 minutes at room temperature. The cells were then washed and incubated with an anti-S protein antibody for 20 minutes at room temperature, followed by staining with a FITC-labeled secondary antibody (111-605-045, Jackson ImmunoResearch). The percentage of FITC-positive cells was determined using a Fortessa X20 flow cytometer (BD Biosciences).

### Detection of FXa binding to S protein-ACE2 complex using ELISA

ACE2 protein was used as a coating reagent in 96-well plates, which were incubated with 1 μg/ml full-length S protein with His tag that had been pretreated with or without FXa (P8010L, NEB) for 2 hours at room temperature. An HRP-conjugated anti-His tag antibody (ab1187, Abcam) was used as a detecting antibody. Absorbance was measured at OD450 nm using a Multiskan™ FC Microplate Photometer (Fisher Scientific).

### Detection of FXa binding to S protein-ACE2 complex using flow cytometry

HEK293T cells stably expressing ACE2 protein were incubated with full-length S protein or FXa-pretreated full-length S protein for 20 minutes at room temperature. Cells were then washed and incubated with an anti-S protein antibody for 20 minutes at room temperature, followed by staining with an APC-labeled secondary antibody (111-005-003, Jackson ImmunoResearch). The percentage of APC-positive cells was determined using a Fortessa X20 flow cytometer (BD Biosciences).

### In vivo infection model

6-8-week-old K18-hACE2 mice were anesthetized with ketamine (80 mg/kg)/xylazine (8 mg/kg) and intranasally infected with 5×10^3^ PFU wild type SARS-CoV-2 or B.1.1.7 variant in 25 μl DMEM, followed by intranasal treatment with PBS, FXa-Fc (200 μg), or Fc (200 μg) in 25 μl DMEM. Infected mice were maintained in BCU isolator cages (Allentown, NJ, USA) in the NAU ABLS3. Mice were then treated with PBS or rivaroxaban (30 mg/kg) via gavage or fondaparinux (30 mg/kg) via intraperitoneal injection for 4 times at a frequency of every other day *(30,31)*. Body weights of mice were monitored daily. Mice were euthanized using ketamine (100 mg/kg)/xylazine (10 mg/kg) when body weights dropped below 20% of their original body weights. RNA was isolated from trachea, lung, and brain tissues to assess viral load using quantitative real-time PCR as described below. Expression of SARS-CoV-2 viral protein NP was examined using immunohistochemistry (IHC) in the trachea, lung, and brain sections from infected mice as described below. Experiments and handling of mice were conducted under federal, state, and local guidelines and with approvals from the Northern Arizona University (20-005) and City of Hope Animal Care and Use Committees.

### Quantitative real-time PCR

Mouse tissues were homogenized in DMEM and RNA was isolated using a PureLink RNA isolation kit (K156002, Invitrogen). Viral copy numbers were detected using the One-Step QPCR kit (1725150, BioRad).

### H&E and IHC

4-μm-thick sections were cut from paraffin blocks of lung and liver tissues from COVID-19 patients and non-COVID-19 donors. IHC staining with an anti-FXa protein antibody (PIPA529118, Invitrogen), an anti-furin antibody (ab183495, Abcam), an anti-trypsin antibody (ab200997, Abcam), or an anti-plasmin antibody (LS-C150813-1, LSBio) as a primary antibody was performed by the Pathology Shared Resource Core at City of Hope Beckman Research Institute. Stained slides were mounted and scanned for observation.

Mouse tissues isolated from experimental mice were placed in 10% neutral buffered formalin for a minimum of 72 hours. After paraffin embedding, 4-μm-thick sections were cut from the blocks. H&E staining and IHC with anti-NP protein antibody (NB100-56576, Novus) as the primary antibody were performed by the Pathology Shared Resource Core at City of Hope Beckman Research Institute. Stained slides were mounted and scanned for observation.

### Statistical analysis

Prism software v.8 (GraphPad, CA, USA) and SAS v.9.4 (SAS Institute. NC, USA) were used to perform statistical analyses. For continuous endpoints that were normally distributed or normally distributed after log transformation such as MFI or copy number, Student’s *t* test or paired *t* test was used to compare two independent or matched groups, respectively. One-way ANOVA models or generalized linear models were used to compare three or more independent groups. For data with repeated measures from the same subject, linear mixed models were used to account for the variance and covariance structure due to repeated measures. Survival functions were estimated using the Kaplan–Meier method and compared using the two-sided log rank test. All tests were two-sided. P values were adjusted for multiple comparisons using Holm’s procedure. A P value of 0.05 or less was considered statistically significant. The p-values are represented as: * <0.05, ** <0.01, *** <0.001, and **** <0.0001

## ACKNOWLEDGMENTS

This work was supported by grants from the NIH (NS106170, AI129582, CA247550, and CA223400 to J. Yu; CA210087, CA068458, and CA163205 to M.A. Caligiuri), the Leukemia and Lymphoma Society (1364-19 to J. Yu), The California Institute for Regenerative Medicine (DISC2COVID19-11947 to J. Yu), and The Flinn Foundation grant 2304 (PK and BB) and Arizona Board of Regents TRIF award (PK).

## AUTHORS’ CONTRIBUTIONS

J. Yu, M.A. Caligiuri, and W. Dong conceived and designed the project. J. Yu, M.A. Caligiuri, W. Dong, J. Wang, L. Tian, H. Mead, S. Jaramillo, E. Settles, P. Keim and B. Barker designed and supervised experiments conducted in the laboratories. Dong, J. Wang, L. Tian, J. Zhang, A. Li, performed experiments and/or data analyses. R. Zumwalt provided the COVID-19 patient samples. S. Whelan provided VSV-SARS-CoV-2. W. Dong, L. Tian, E. Settles, P. Keim, B. Barker, J. Yu, and M.A. Caligiuri wrote, reviewed and/or revised the paper. All authors discussed the results and commented on the manuscript.

