## supplemental data for "FXa cleaves the SARS-CoV-2 spike protein and blocks cell entry to protect against infection with inferior effects in B.1.1.7 variant"

### EXTENDED DATA FIGURES and LEGENDS:

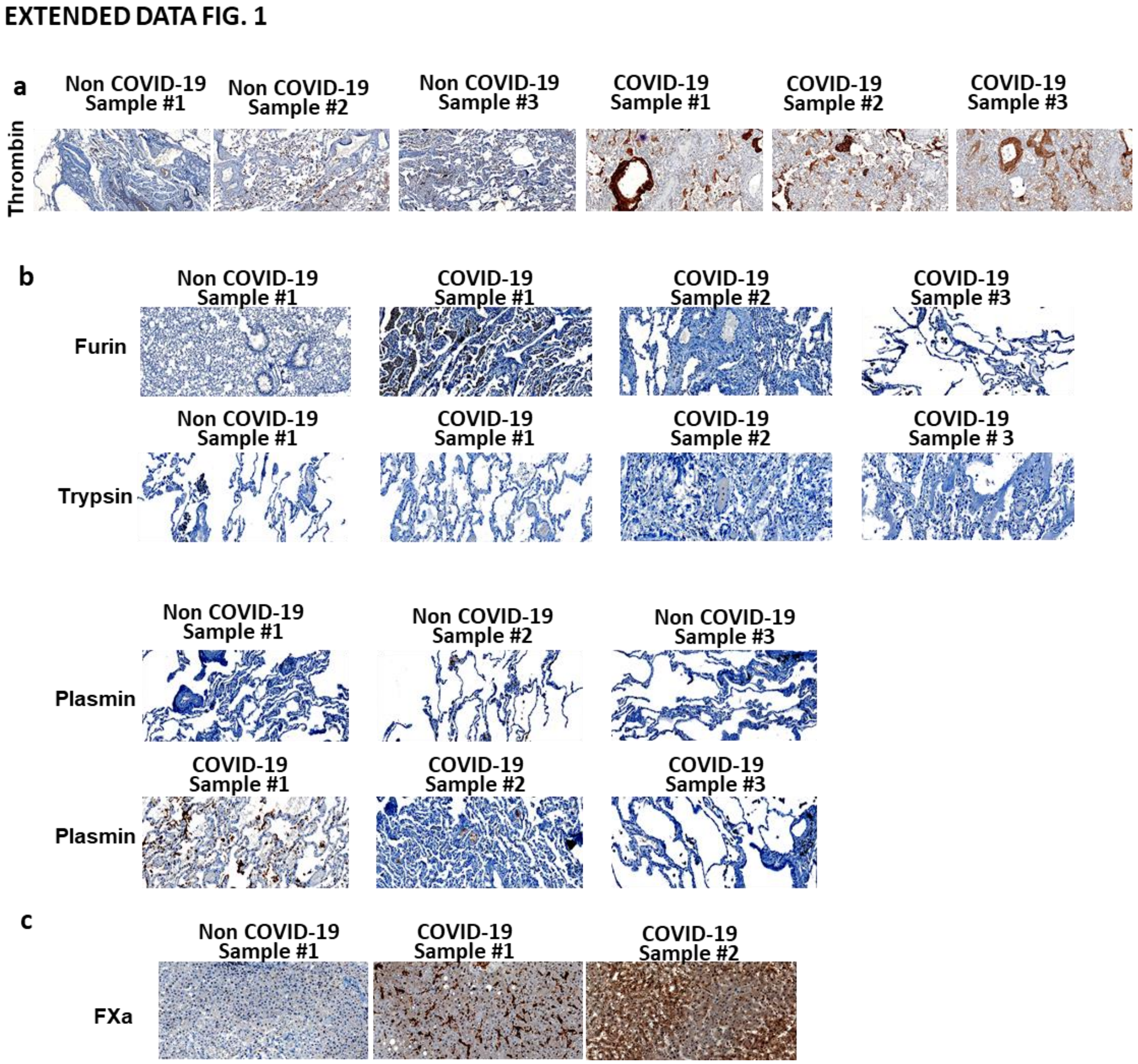


**Extended Data Fig 1. Expression of SP in organs of autopsy samples of patients who died of COVID-19 vs. non-COVID-19 donors**. (**a**) Expression of thrombin in the lung of autopsy samples of patients who died of COVID-19 vs. non-COVID-19 donors. **(b)** Expression of furin, trypsin, and plasmin SPs in the lung of autopsy samples of patients who died of COVID-19 vs. non-COVID-19 donors. **(c)** Expression of FX SP in the liver of autopsy samples of patients who died of COVID-19 vs. non-COVID-19 donors.

**EXTENDED DATA FIG. 2**

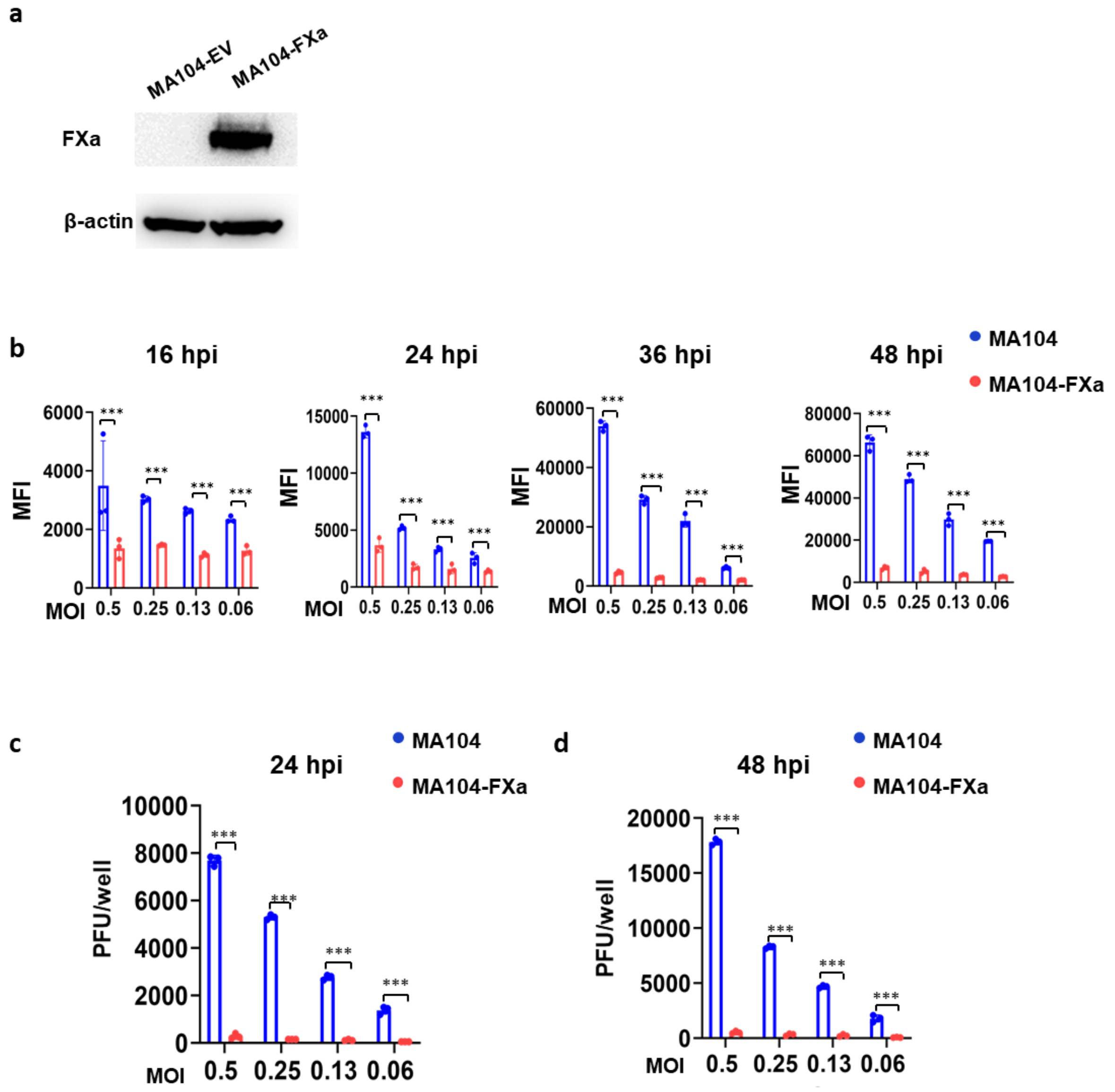


**Extended Data Fig 2. Infectivity and virus production of VSV-SARS-CoV-2 in MA104 cells expressing FXa or a control vector.** (**a**) Confirmation of forced over-expression of FXa in MA104 cells was conducted by immunoblotting. (**b**). MA104 cells transduced with FXa (MA104-FXa) or an empty vector (MA104-EV) were infected by VSV-SARS-CoV-2. Infectivity of the cells were quantified by flow cytometry at 16, 24, 36 and 48 hpi. (**c and d**) The titer of the supernatant of VSV-SARS-CoV-2-infected MA104 or -MA104-FXa cells at 24 hpi and 48 hpi was determined by re-infection of Vero cells. All data are representative of at least three independent experiments. Experiments in A is representative of three independent experiments with similar data. For all panels, error bars indicate SD, and statistical analyses were performed by means of Student's t tests. ***P ≤ 0.001.

**EXTENDED DATA FIG. 3**

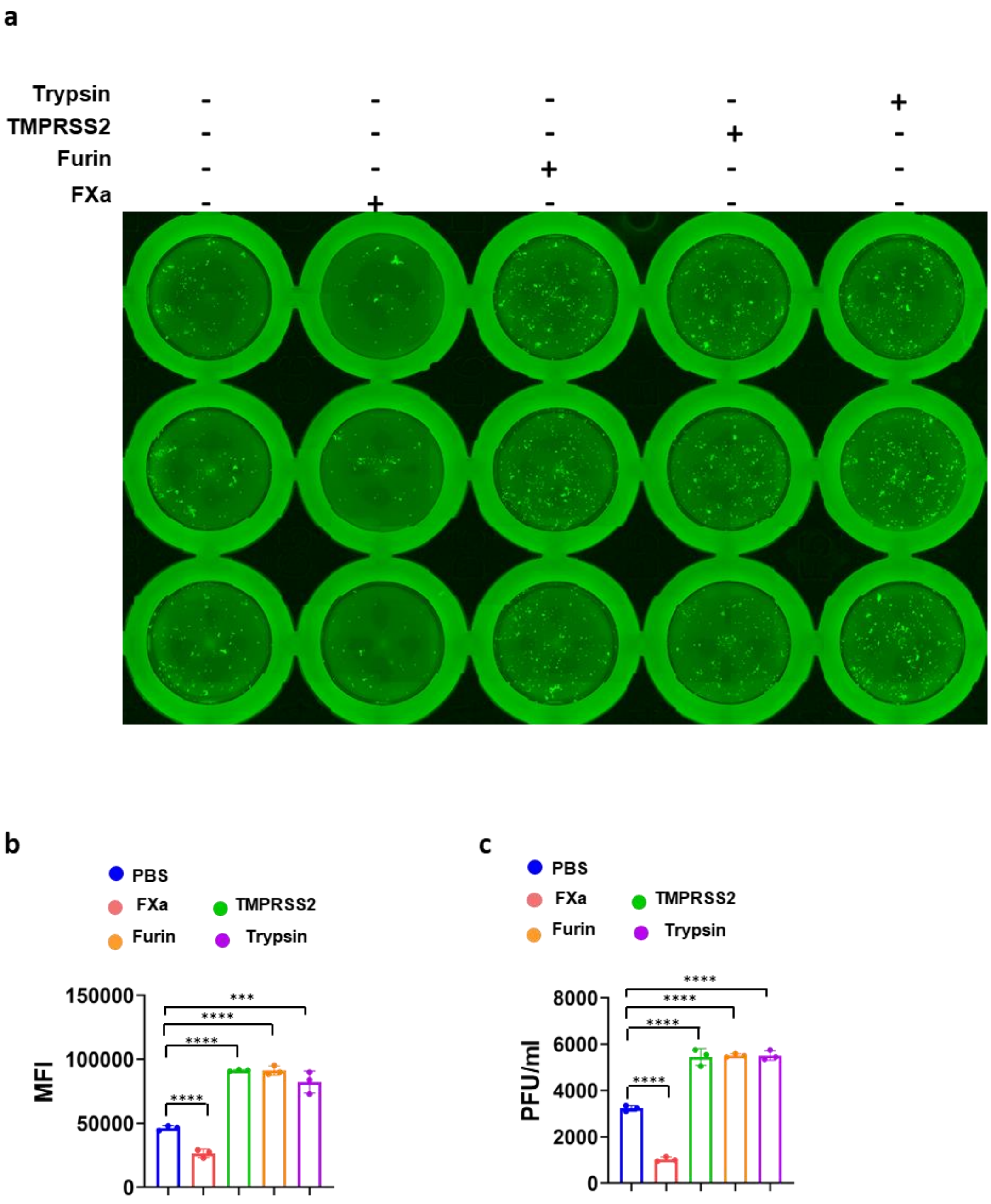

**Extended Data Fig 3**. **FXa inhibits while TMPRSS2, trypsin, and furin promote VSV-SARS-CoV-2 infection in MA104 cells.** (**a and b**) Infectivity of MA104 cells infected with VSV-SARS-CoV-2 in the presence or absence of FXa, TMPRSS2, trypsin, or furin was determined by fluorescence microscope. (**c**) Virus production MA104 cells infected with VSV-SARS-CoV-2 in the presence or absence of FXa, TMPRSS2, trypsin, or furin was determined by re-infection of Vero cells. All data are representative of at least three independent experiments. Experiments in A is representative of three independent experiments with similar data. For all panels, error bars indicate SD, and statistical analyses were performed by one-way ANOVA models. **$P \leq 0.01$; n.s, not significant.

EXTENDED DATA FIG. 4

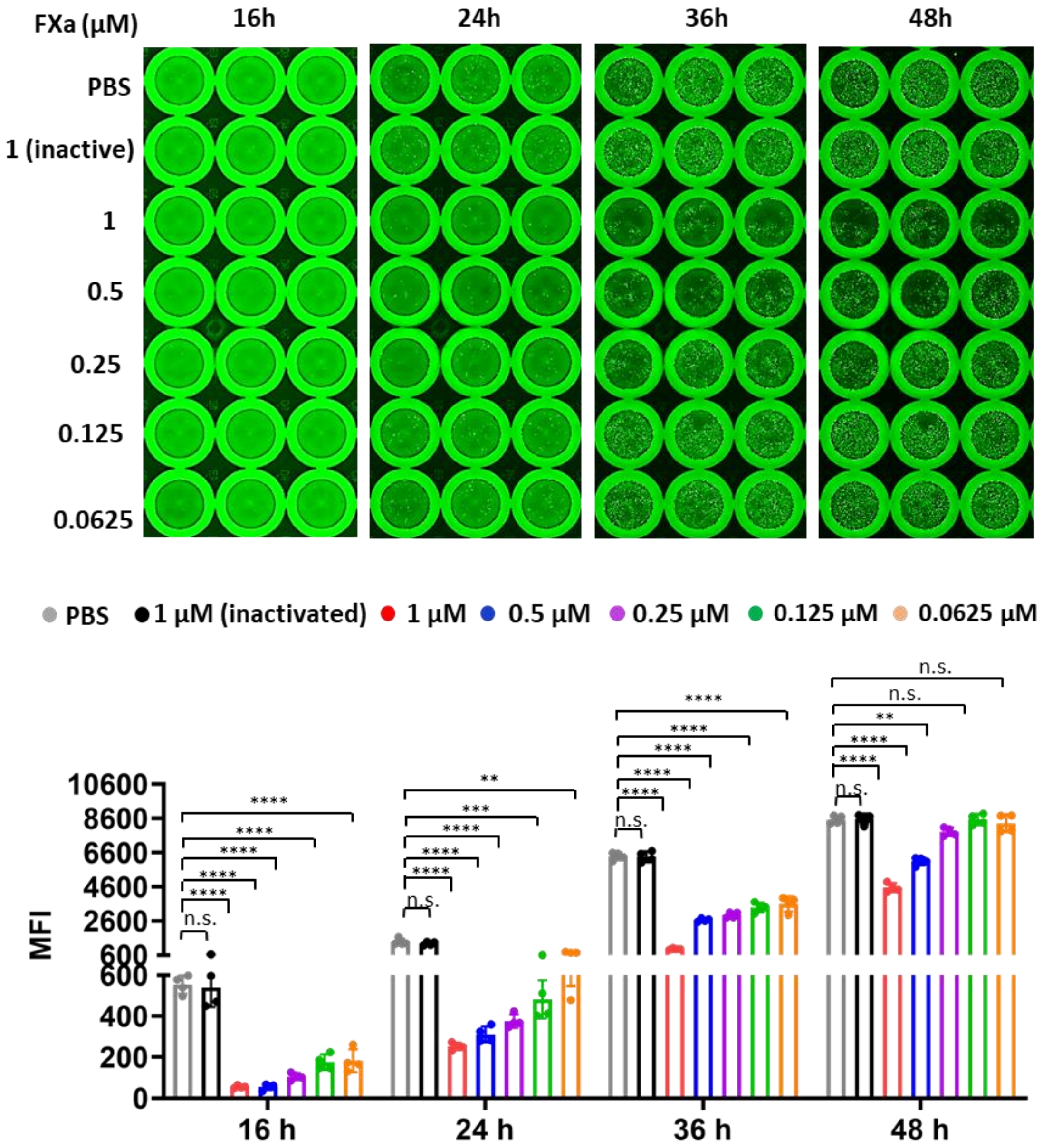


**Extended Data Fig 4. FXa inhibits VSV-SARS-CoV-2 infection.** VSV-SARS-CoV-2 was preincubated with FXa at different concentrations 1 hour before infection. Cells were imaged at 16, 24, 36 and 48 hpi by fluorescence microscopy (up panel) and the corresponding infectivity was measured by flow cytometry (bottom panel). Error bars indicate SD, and statistical analyses were performed by one-way ANOVA models. ***P ≤ 0.001; n.s, not significant.

**EXTENDED DATA FIG.5**

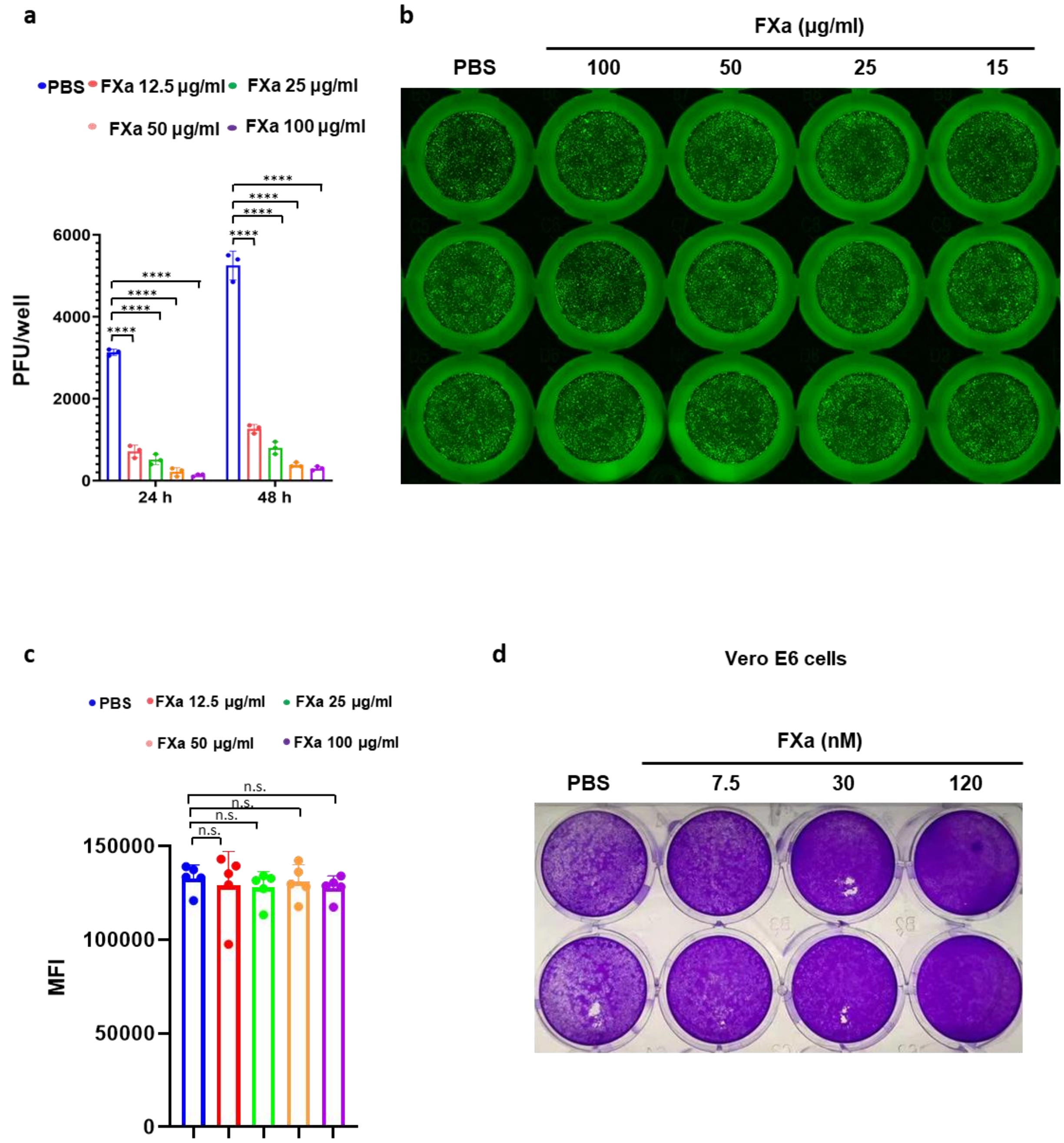


**Extended Data Fig 5. Determination of the effect of FXa on virus particles and host cells.** (**a**) VSV-SARS-CoV-2 was preincubated with FXa at different concentrations 1 hour prior to viral infection of MA104 cells. The supernatants were collected at 24 and 48 hpi for virus production assay by re-infection of Vero cells. (**b and c**). MA104 cells were preincubated with or without FXa at different concentrations 1 hour before infection. Infectivity of VSV-SARS-CoV-2 in the preincubated or untreated MA104 with FXa was determined by fluorescence microscope (b) and flow cytometry (c). **(d)** Vero E6 cells were infected with live wild-type SARS-CoV-2. At 24 hpi, infectivity was measured by a traditional plaque assay. All data are representative of at least three independent experiments. Experiments in b are representative of three independent experiments with similar data. For all panels, error bars indicate SD, and statistical analyses were performed by one-way ANOVA models. *P ≤ 0.05; **P ≤ 0.01; ****P ≤ 0.0001; n.s, not significant.

### EXTENDED DATA FIG. 6

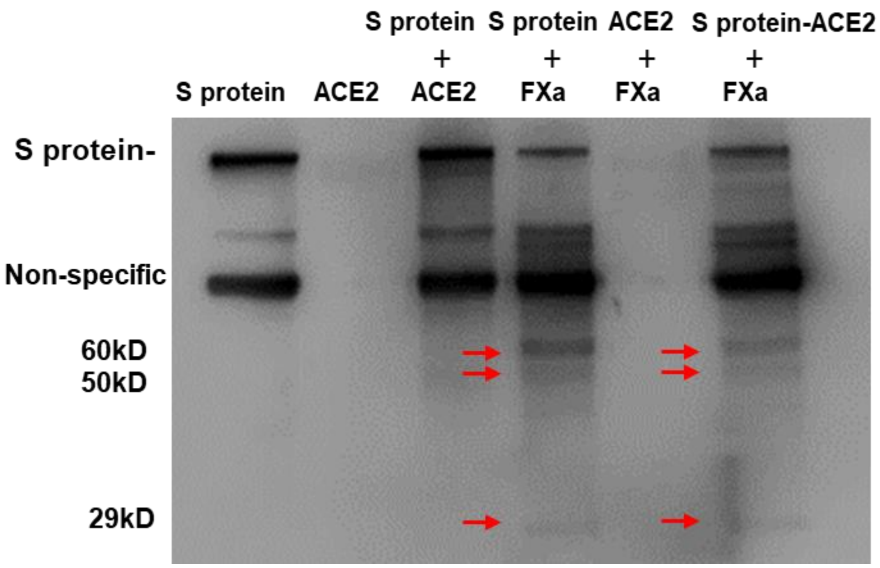


**Extended Data Fig 6. Cleavage of S protein in the S protein-ACE2 complex by FXa.** S protein was pre-incubated with ACE2 for 1 hour, followed by addition of FXa for another 1 hr. The cleavage of S protein in the S protein-ACE2 complex was determined by immunoblot. The experiment is representative of at least three independent experiments.

**EXTENDED DATA FIG. 7**

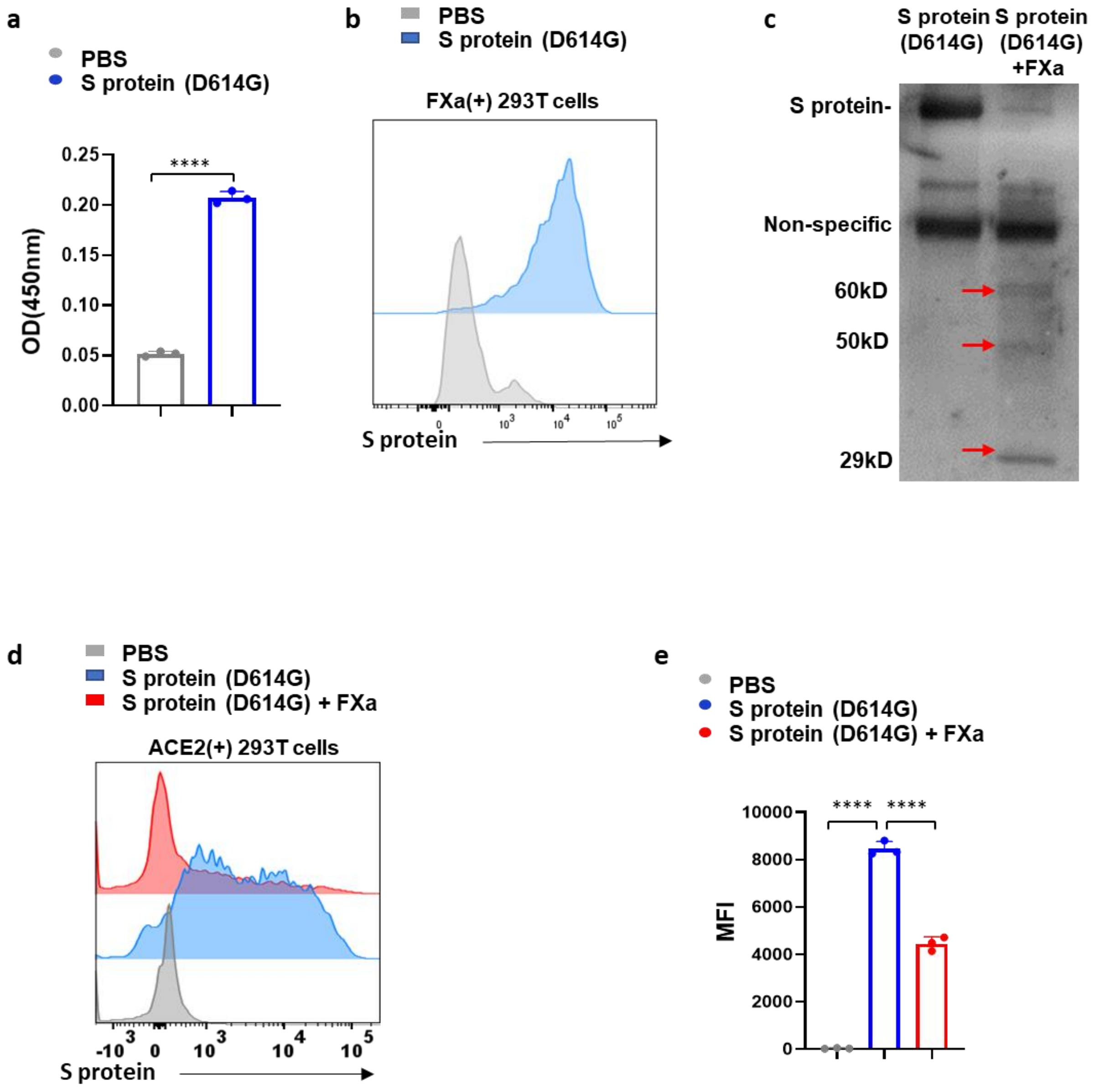


**Extended Data Fig 7. Binding of FXa with and cleavage of the mutant S protein of the SARS-CoV-2 D614G variant.** (**a**) The binding affinity of FXa and D614G S protein was measured by ELISA. (**b**) The binding of the mutant S protein of the SARS-CoV-2 D614G variant with FXa expressed on 293T cells was assessed by flow cytometry. (**c**) Cleavage assay of D614G S protein by FXa was measured by immunoblotting. (**d and e**) The binding affinity of ACE2 and D614G S protein pretreated with or without FXa was measured by flow cytometry. All data are representative of at least three independent experiments. Experiments in c is representative of three independent experiments with similar data.  For all panels, error bars indicate SD, and statistical analyses were performed Student's t test (a) and one-way ANOVA models (e). ****P ≤ 0.0001; n.s, not significant.

### EXTENDED DATA FIG. 8

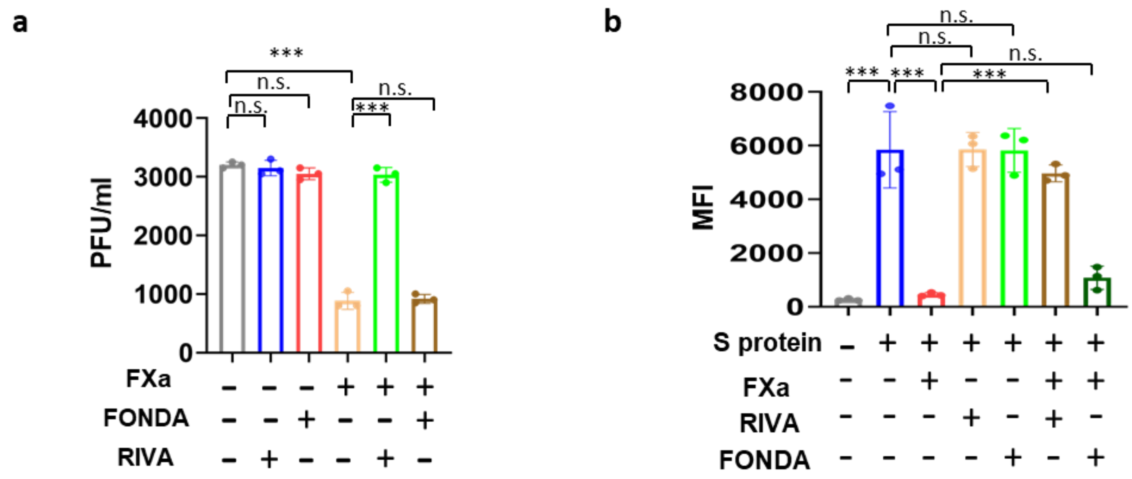


**Extended Data Fig 8. The effect of RIVA or FONDA on infectivity and virus production of VSV-SARS-CoV-2 or live SARS-CoV-2 pretreated with FXa**. (**a**) Virus production of FXa-pretreated vs. untreated VSV-SARS-CoV-2 in MA104 cells in the presence or absence of RIVA or FONDA was determined by re-infection of Vero cells. (**b**) FXa pretreated with or without RIVA or FONDA was incubated with S protein, followed by assessing the binding capability of these S proteins binding with ACE2 expressed on 293T cells by flow cytometry (summary data of main Fig. 3l). All data are representative of at least three independent experiments. For all panels, error bars indicate SD, and statistical analyses were performed by one-way ANOVA models. ***P ≤ 0.001; n.s, not significant.

EXTENDED DATA FIG. 9

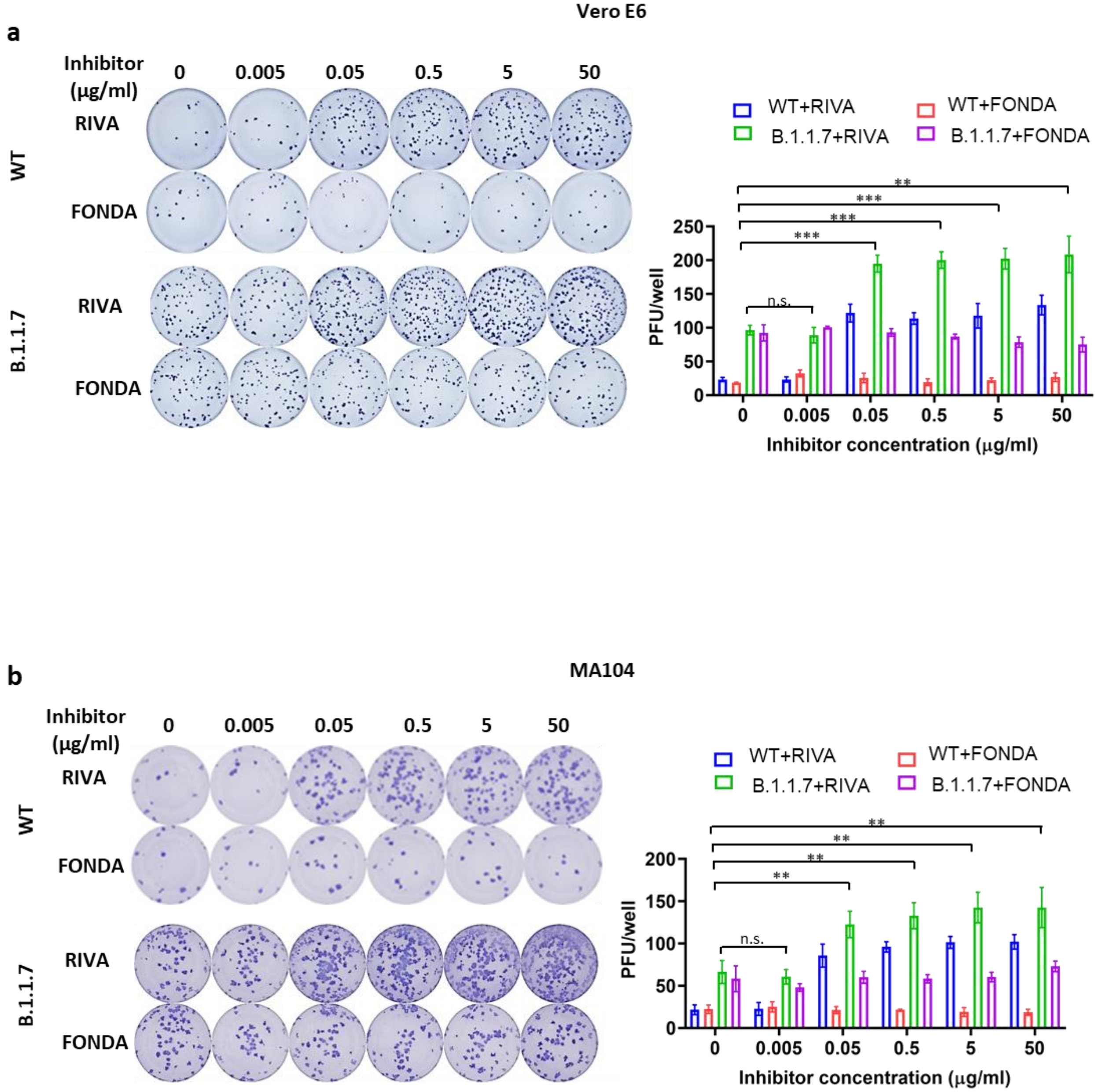


**Extended Data Fig 9. The effect of various doses of RIVA or FONDA on infectivity of live wild-type SARS-CoV-2 or the B.1.1.7 variant pretreated with FXa in Vero E6 and MA104 cells**. **(a and b)** Vero E6 (a) and MA104 (b) cells were infected with live wild-type SARS-CoV-2 or the B.1.1.7 variant pretreated with different doses of RIVA or FONDA in the presence of FXa. At 24 hpi, viral infectivity was measured by immune-plaque assay. A representative assay is presented on the left at the summary data on the right. For all panels, error bars indicate SD, and statistical analyses were performed by one-way ANOVA models. ***P ≤ 0.001; n.s, not significant.

EXTENDED DATA FIG. 10

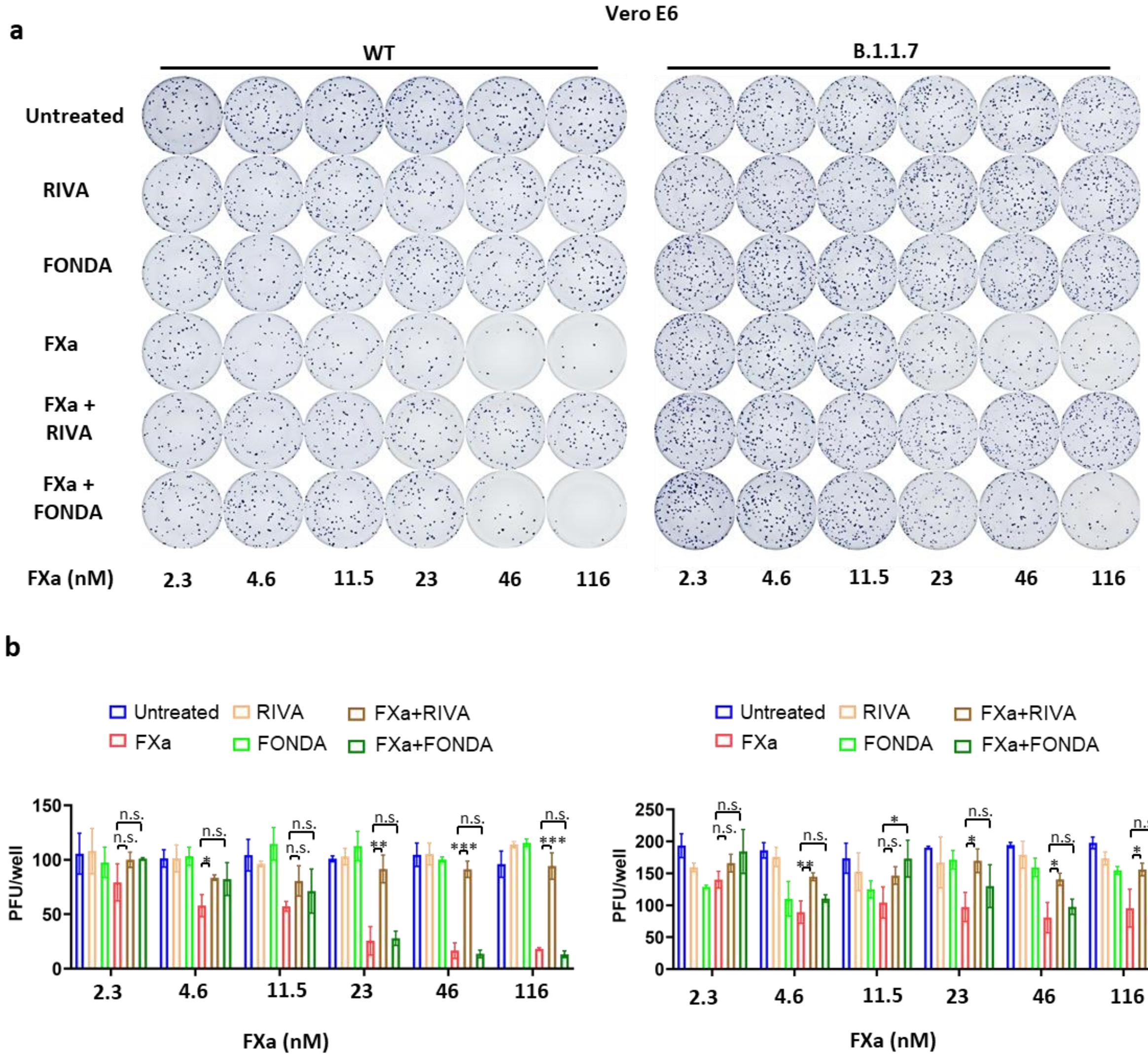


**Extended Data Fig 10. The effect of RIVA or FONDA on infectivity of live wild-type SARS-CoV-2 or the B.1.1.7 variant pretreated with various doses of FXa in Vero E6 cells**. **(a and b)** Vero E6 cells were infected with live wild-type SARS-CoV-2 or the B.1.1.7 variant pretreated with different doses of FXa in the presence of RIVA or FONDA. At 24 hpi, viral infectivity was measured by immune-plaque assay (a). (b) The summary data of (a). For all panels, error bars indicate SD, and statistical analyses were performed by one-way ANOVA models. ***P ≤ 0.001; n.s, not significant.

**EXTENDED DATA FIG. 11**

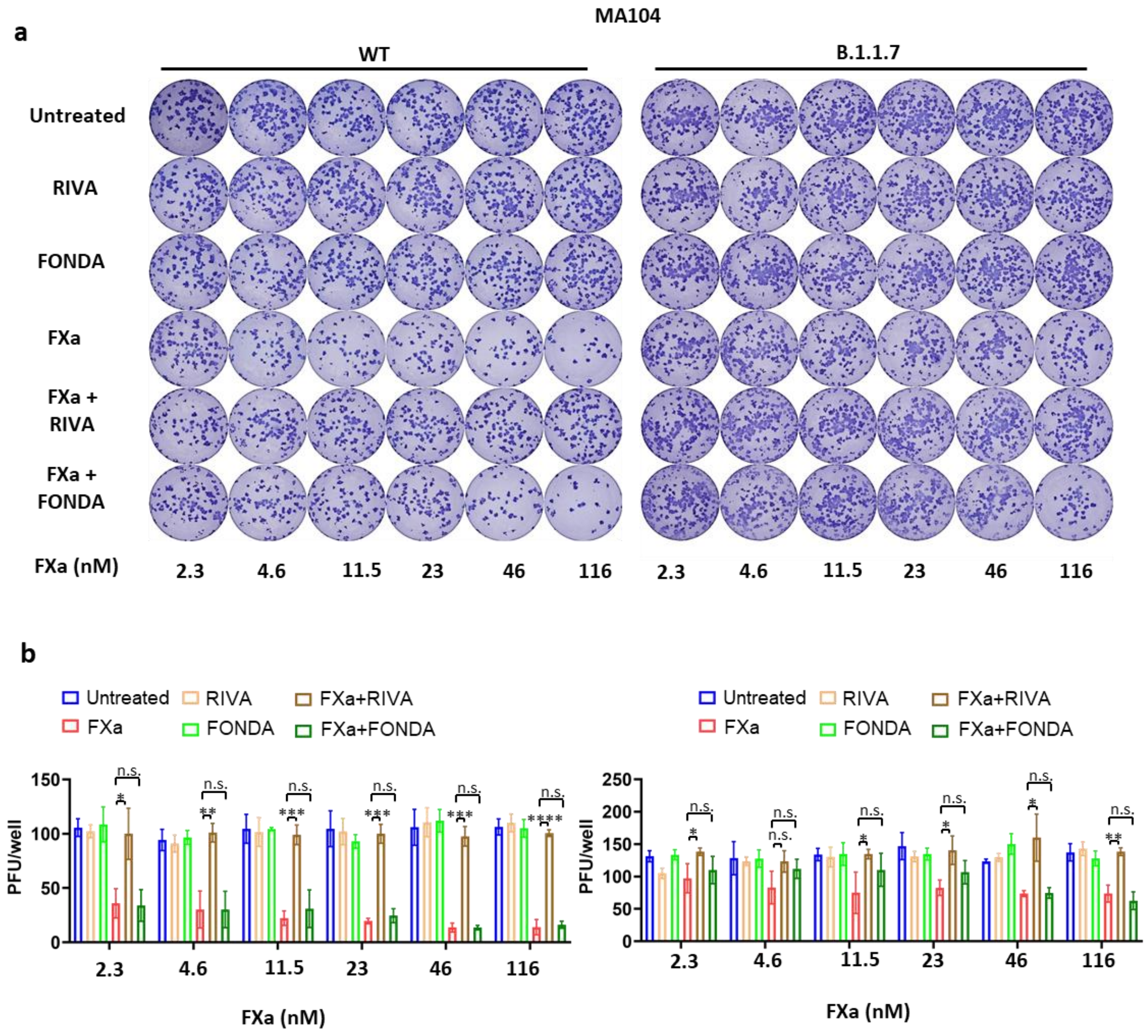


**Extended Data Fig 11. The effect of RIVA or FONDA on infectivity of live wild-type SARS-CoV-2 or the B.1.1.7 variant pretreated with various doses of FXa in MA104 cells**. **(a and b)** MA104 cells were infected with live wild-type SARS-CoV-2 or the B.1.1.7 variant pretreated with different doses of FXa in the presence of RIVA or FONDA. At 24 hpi, viral infectivity was measured by immune-plaque assay (a). (b) The summary data of (a). For all panels, error bars indicate SD, and statistical analyses were performed by one-way ANOVA models. ***$P \leq 0.001$; n.s, not significant.
